# PRDM16 establishes lineage-specific transcriptional program to promote temporal progression of neural progenitors in the mouse neocortex

**DOI:** 10.1101/573857

**Authors:** Li He, Jennifer Jones, Weiguo He, Bryan Bjork, Jiayu Wen, Qi Dai

**Author notes:** Equal contribution (Li He, Jennifer Jones). Co-senior authors (Bryan Bjork, Jiayu Wen, Qi Dai).

## Abstract

Radial glia (RG) in the neocortex sequentially generate distinct subtypes of projection neurons, accounting for the diversity and complex assembly of cortical neural circuits. Mechanisms that drive the rapid and precise temporal progression of RG are beginning to be elucidated. Here we reveal that the RG-specific transcriptional regulator PRDM16 promotes the transition of early to late phases of neurogenesis in the mouse neocortex. *Prdm16* mutant RG delays the timely progression of RG, leading to defective cortical laminar organization. We show that PRDM16 regulates expression of neuronal specification genes and a subset of genes that are dynamically expressed between mid-and late-neurogenesis. Our genomic analysis suggests that PRDM16 suppresses target gene expression through maintaining chromatin accessibility of permissive enhancers. Altogether, our results demonstrate a critical role of PRDM16 in establishing stage-specific gene expression program of RG during cortical neurogenesis. These findings also support a model where progenitor cells are primed with daughter cell gene expression program for rapid cell differentiation.

## Introduction

Radial glia in the developing mammalian cerebral cortex are neural stem cells that give rise to all excitatory neurons (Anthony et al. 2004; Kriegstein and Alvarez-Buylla 2009). During peak phase of neurogenesis, RG divide asymmetrically to produce a self-renewing RG and a neuron or a transit-amplifying intermediate progenitor (IP) that divides again to produce more neurons (Noctor et al. 2004). On each embryonic (E) day starting at E11.5, RG generate a new laminar layer with distinct neuronal subtypes. Layer (L) 6 neurons are born first at E12.5, followed by L5 (E13.5), L4 (E14.5) and L2/3 (E15.5) neurons. The newborn neurons migrate along the radial fiber of their mother RG, passing through and positioning on top of earlier-born neurons (Angevine and Sidman 1961; Okano and Temple 2009; Kwan et al. 2012). Thus, the identity and laminar position of neuronal subtypes are tightly linked to their birthdate. During developmental progression, the competence of progenitors becomes progressively restricted (Frantz and McConnell 1996; Desai and McConnell 2000; Gaspard et al. 2008; Gao et al. 2014). Previous studies have suggested that both extrinsic and intrinsic mechanisms are needed to control temporal identity of neural progenitors (McConnell and Kaznowski 1991; Chenn and Walsh 2002; Fukumitsu et al. 2006; Ge et al. 2006; Shen et al. 2006; Hsu et al. 2015; Dennis et al. 2017; Zahr et al. 2018).

In differentiating neurons, cell-specific transcription factors and their regulated transcriptional cascades further guide neuronal specification, migration and circuit assembly(Greig et al. 2013). For example, complex interplay between the deep layer factor Tbr1, mid-layer Fezf2 and upper-layer Satb2 guide specification of corticothalamic, subcerebral and callosal neurons in deep-, mid-and upper-cortical layers (Srinivasan et al. 2012; McKenna et al. 2015). Two related POU domain transcription factors, Pou3f2 (Brn2) and Pou3f3 (Brn1), are also required for determining the identity and migration of upper layer neurons (McEvilly et al. 2002; Sugitani et al. 2002). The proteins of these factors serve as subtype-specific markers (Molyneaux et al. 2007). Their mRNAs, as suggested by a few recent studies, exist in RG in much earlier stages (Yoon et al. 2017; Zahr et al. 2018). It is an interesting question whether the presence of the neuronal gene mRNAs in RG is important for RG neurogenesis.

The choroid plexus (ChP) protects the brain via the blood-CSF (cerebrospinal fluid) barrier and regulates CSF composition via specific and regulated transfer and secretion(Lehtinen et al. 2013; Johansson 2014). Signaling molecules in CSF (e.g. Shh, Igf1, Wnt4, Tgm2 and Fgf2) are delivered to NSCs and directly influence NSC behavior (Imayoshi et al. 2008; Lehtinen et al. 2011; Johansson et al. 2013; Johansson 2014). However, mechanisms and factors controlling development of the ChP are not fully understood.

The PR domain-containing (PRDM) family protein PRDM16 is a key transcriptional regulator in diverse cell types (Kajimura et al. 2008; Chuikov et al. 2010; Aguilo et al. 2011). In embryonic and adult brain, PRDM16 was shown to control neural stem cell maintenance(Chuikov et al. 2010; Shimada et al. 2017), IP proliferation (Baizabal et al. 2018), neuronal migration (Inoue et al. 2017; Baizabal et al. 2018) and ependymal cell differentiation (Shimada et al. 2017). It was shown that in these contexts PRDM16 regulates genes involved in reactive oxygen species (ROS) levels (Chuikov et al. 2010; Inoue et al. 2017) and epigenetic states of its bound enhancers (Baizabal et al. 2018). The PRDM16 protein (**Supplemental Fig. S1A**) contains a PR domain that possesses intrinsic Histone H3K4 (Zhou et al. 2016) and H3K9 methyltransferase activity (Pinheiro et al. 2012), two potential DNA binding zinc-finger clusters (Nishikata et al. 2003) and interaction motifs for the co-repressors CtBP1/2. The transcriptional activity of PRDM16 is context-dependent (reviewed in (Chi and Cohen 2016)), as it activates gene expression when associated with other activators and represses gene expression when interacting with co-repressors.

In this study, we explored the *in vivo* function of *Prdm16* in the developing mouse brain. We show that *Prdm16* controls brain development in at least two areas, the ChP and the neocortex. *Prdm16* is essential for ChP development. In the neocortex, *Prdm16* promotes the shift between L5 neuron and L2-4 neuron specification. PRDM16 sets up the transcriptional landscape for mid-layer and upper-layer specification genes and influences gene expression dynamics of RG between mid-and late-neurogenesis. Together, our findings suggest that the gene expression program established by PRDM16 confers temporal identity of RG at the onset of early and late neurogenesis transition.

## Results

### Prdm16 is required for neocortical development and choroid plexus formation

To assess the function of *Prdm16* in the developing brain, we made use of three multifunctional conditional gene trap (cGT) alleles (Strassman et al. 2017) (**Supplemental Fig. 1B**). The *Prdm16^cGT^* (*cGT*) and *Prdm16^cGTreinv^* (*cGTreinv*) mouse strains produce a null allele (**Supplemental Fig.S1B**) and will be referred as *Prdm16* KO mutants. To examine PRDM16 activity in the neocortex, we depleted *Prdm16* expression in the forebrain using the *Emx1^tm1(cre)Krj/J^* (*Emx1^IREScre^*) deleter strain (Gorski et al. 2002) and the conditional *Prdm16^cGTinv^* (*cGTinv*) strain. *cGTinv* will be referred to as *Prdm16* cKO throughout this manuscript. The *Prdm16* transcript is detectable in E9.5 brain (**Supplemental Fig.S1C** and (Horn et al. 2011)). At E13.5, PRDM16 has specific expression in the ChP and in the ventricular zone (VZ) where it co-localizes with the RG marker Sox2 (**Supplemental Fig.S1D**). In KO animals, PRDM16 staining is lost in the entire brain (**Supplemental Fig. S1D**), while in cKO mutants it is depleted in the dorsal telencephalon but remains expressed in the ventral telencephalon and the ChP (**Supplemental Fig. S1E**).

**Figure 1.**
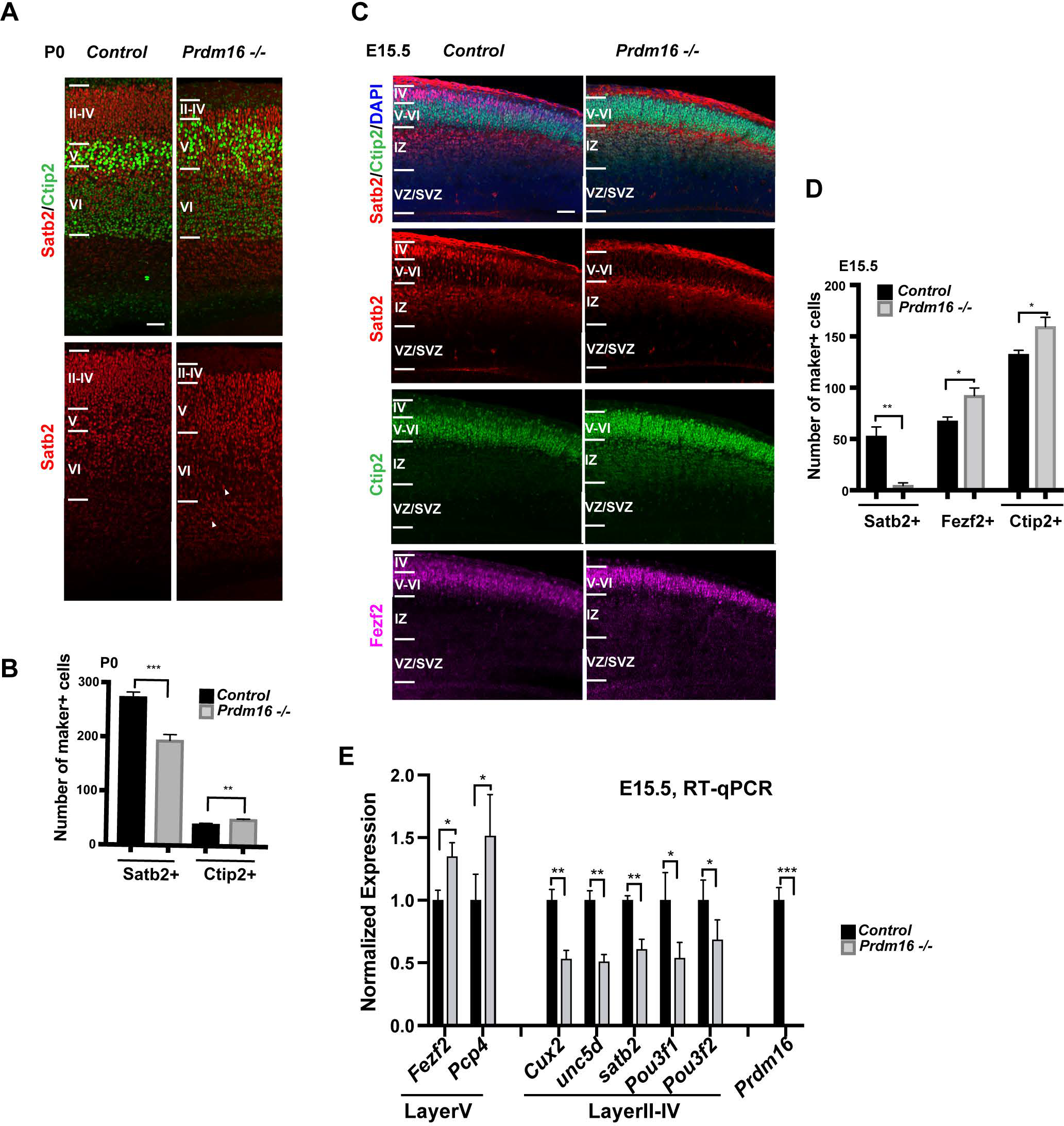
PRDMl6 regulates cortical laminar organization. **(A)** Images of P0 cortices show increased Ctip2+ mid-layer and reduced Satb2+ upper-layer. White arrow heads indicate retained Satb2+ in the lower layers. Cortical layers are highlighted according to relative distribution of Satb2+ and Ctip2+ cells. **(B)** Quantification of the marker+ cells in (A) in 80 µm column across the P0 cortex (n=3). **(C)** Images of El5.5 cortices show reduction of the Satb2+ layer and expansion of the Ctip2+ or Fezf2+ layer. **(D)** Quantification of the marker+ cells in (C) in I 00 µm column across the cortex of El 5.5 (n=3). **(E)** Measurement of layer marker genes by RT-qPCR from El 5.5 control and Prdml6 KO cortices. All data are shown as mean +/−SD; *p<0.05; **p<0.01; ***p<0.001. Scale bar: 50 µm.

We first analyzed the cortical laminar organization of the KO brains, by labeling cortical neurons with Satb2 for the upper-layer (L2-4, II-IV) and Ctip2 and Fezf2 for the mid-layer n (L5, V, strong Ctip2 and Fezf2) and the deep-layer (L6, VI, weak Ctip2 and Fezf2). At postnatal day 0 (P0), mutant cortices showed expansion of Ctip2+ layer, accompanied by thinning of Satb2+ upper-layer (**Fig. 1A–B**), compared with control cortices. Some Satb2+ neurons scattered inside the deep layer, suggesting that a subset of upper-layer neurons may have failed to migrate. Similarly at E15.5 when upper-layer neurons were just born, the number of Satb2+ neurons was already reduced and the mid-layer neurons labeled with Fezf2 and Ctip2 were expanded in the mutant (**Fig. 1C–D**). The reciprocal changes of L5 and L2-4 marker genes were confirmed by reverse transcription followed by quantitative PCR (RT-qPCR). The levels of the two L5 genes increased to about 150%, while those of the L2-4 genes decreased to 50-70% (**Fig. 1E)**, indicating that gain of mid-layer neurons roughly compensates for loss of upper-layer neurons at E15.5. Hence, the *Prdm16* KO cortex display two types of defects: over production of mid-layer neurons; compromised neuronal production and defective migration of upper layer neurons.

In *Prdm16* KO brains, the prospective ChP in the lateral and the 3^rd^ ventricles are dramatically reduced (**Supplemental Fig. S1D, F, G**, (Bjork et al. 2010; Strassman et al. 2017)), pointing to an essential role of PRDM16 in the ChP development. Together, the phenotypic analyses in *Prdm16* KO mutant indicate that PRDM16 controls brain development in at least two brain areas, the neocortex and the ChP.

### Expression of Prdm16 in the forebrain is responsible for the effects on laminar organization

To test the direct roles of PRDM16 in cortical development, we analyzed *Emx1^IREScre^*-mediated *Prdm16* cKO mutants where *Prdm16* is depleted in the forebrain (**Supplemental Fig. S1E**). The *Prdm16* cKO animals survive to adulthood, allowing examination of postnatal stages. At E15.5. *Prdm16* cKO cortices displayed defects similar to *Prdm16* KO embryos, evidenced by the increase in number of Ctip2+ and Fezf2+ neurons and the reduction in number of Satb2+ neurons (**Fig. 2A-B**). At P15, the cKO cortex showed similar defects on the upper- and mid-layer neurons (**Fig. 2C-E**). The Tbr1-labeled deep-layer is unchanged. Some Satb2+ neurons failed to migrate to the upper-layer and were retained below the cortex, as a chunk of grey matter cells (Heterotopia) (**Fig. 2C-D**). Thus, the forebrain depletion of *Prdm16* led to the same effects on cortical laminar organization as the null KO did: reciprocal changes of L5 and L2-4 neurons and failure of upper-layer neuron migration. This result confirms that the laminar organization phenotypes in the mutant cortex are due to loss of *Prdm16* in the forebrain.

**Figure 2.**
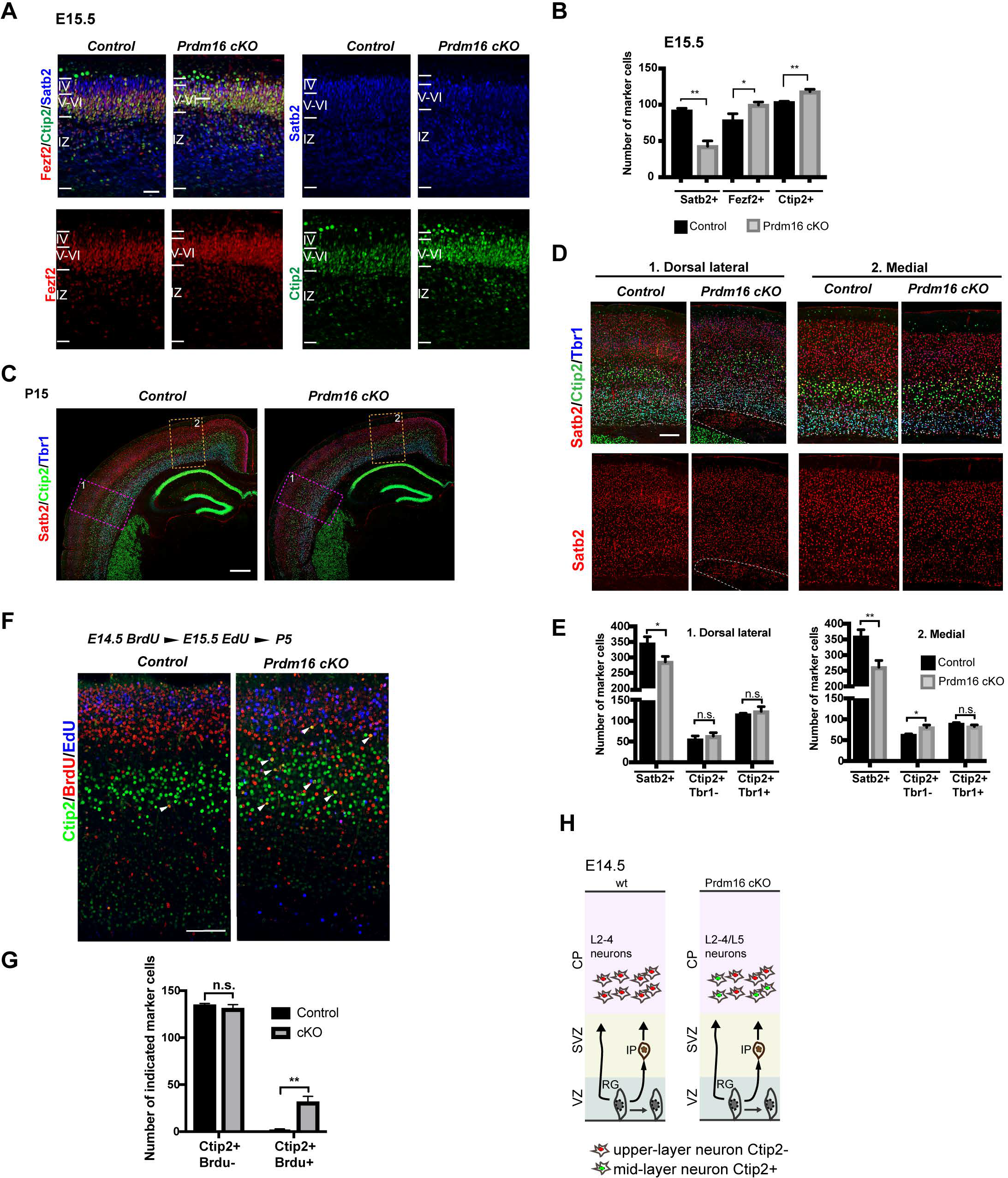
Forebrain-specific depletion of Prdm16 delayed mid-to-late neurogenesis transition. **(A)** Images of E15.5 control and Emx1-Cre:: Prdm16 cKO cortices show reduction of the Satb2+ layer and expansion of the Ctip2+ or Fezf2+ layer. Scale bar: 50 µm. **(B**) Quantification of the marker+ cells in 100 uM column across the cortex (n=3). **(C)** Images of P15 control and Prdm16 cKO cortices stained with Satb2, Ctip2 and Tbr1. Dorsal lateral and medial areas are highlighted in pink and orange rectangles respectively. Scale bar: 100 µm. **(D)** Higher magnification images from C show reciprocal effects on Ctip2+ and Satb2+ layers in cKO cortices and the heterotopia tissue highlighted by white dashed line in the Dorsal lateral region. Scar bar: 50 µm. **(E)** Quantification of the numbers of three cell types in 300 µm column in each area (n=3). **(F)** Images of the P5 control and Prdm16 cKO cortices, stained with Ctip2, BrdU and EdU. White arrowheads point to BrdU+Ctip2+ cells. Scale bar: 100 µm. **(G)** Quantification of the numbers of the Ctip2+BrdU-and Ctip2+BrdU+ cells in 300 µm width across the cortex (n=3). All data are shown as mean +/−SD; *p<0.05; **p<0.01; ***p<0.001. (H) Illustration of the progression delay: mutant RG produce Ctip2+ neurons at E14.5.

### PRDM16 regulates the transition of mid-to-late neurogenesis

We sought to understand the causes of *Prdm16* mutant phenotypes. Given that PRDM16 is a RG-specific factor, PRDM16 may control neurogenesis through modulating intrinsic properties of RG. We reasoned that two possibilities could lead to increase of L5 neurons and decrease of L2-4 neurons. First, if loss of *Prdm16* delayed the transition of neurogenesis from E13.5 to E14.5, mutant RG would produce L5 neurons even after E13.5, which could result in fewer L2-4 neurons. Second, if loss of *Prdm16* increased proliferation of RG at E13.5 and reduced it at E14.5 and later, more daughter neurons could be produced at E13.5 and fewer produced at later time.

To test if PRDM16 controls the timing of RG transition, we traced RG daughter cell fate by injecting pregnant mice with BrdU at E14.5 and EdU at E15.5, and examined the distribution of BrdU and EdU cells and their cellular identity in the P5 cortex. Ctip2+ L5 neurons are born at E13.5 and should not be labeled with BrdU or EdU (Desai and McConnell 2000; Gaspard et al. 2008). As expected, in the control cortex the Ctip2+ cells were rarely labeled with BrdU or EdU (**Fig. 2F–G**). In contrast, the mutant cortex appeared supernumerary Ctip2+BrdU+ neurons, suggesting Ctip2+ neurons were produced even at or after E14.5 in the mutant. Notably, the numbers of the Ctip2+BrdU-cells did not differ between control and mutant, indicating that production of Ctip2+ neurons before E14.5 was normal in the mutant. These results demonstrate that some of *Prdm16* mutant RG cells failed to transit from E13.5 to E14.5 and continued to produce Ctip2+ neurons at E14.5 (**Fig. 2H**).

BrdU+ or EdU+ cells were also found in the heterotopia and the deep layer (**Supplemental Fig. S2A-C**), confirming that the retained cells were the upper-layer neurons that failed to migrate but not from cell-fate transformation in the deep layer or heterotopia. None of the Ctip2+BrdU+ cells were retained in the deep layer (**Fig. 2F**) or in the heterotopia (**Supplemental Fig. S2B**), suggesting that even the latter-produced Ctip2+ neurons migrate normally and that the migration failure is specific to upper-layer neurons.

### PRDM16 promotes proliferation of intermediate progenitors during late neurogenesis

To test if PRDM16 regulates proliferation, we examined RG and IP cell counts at E15.5 by labeling RG with Pax6 and IPs with Tbr2. Remarkably, there was a reduction in the number of Tbr2+ IPs in the cKO cortex, whereas Pax6+ RG were not affected (**Fig. 3A-B**). We further assessed proliferation of IP cells by EdU labeling. We injected EdU to pregnant mice with embryos at E15 and analyzed the brains after 12 hours. There was a significant increase of the percentage of Edu+Ki67-cells over all EdU+ cells, indicating more cells exiting cell cycle in mutant (**Fig. 3C-D**). We observed a significant decrease of Ki67+ cells specifically in the mutant SVZ (**Fig. 3C, 3E**), suggesting decreased proliferation of IP cells. To confirm this, we injected animals with EdU at E15.5 and waited for 2 hours before harvesting. The fraction of EdU+Tbr2+ cells over all EdU+ cells is significantly less in mutant compared with control (**Fig. 3F-G**), indicating that mutant cortex had fewer Tbr2+ cells in S phase presumably due to fewer proliferative IPs.

**Figure 3.**
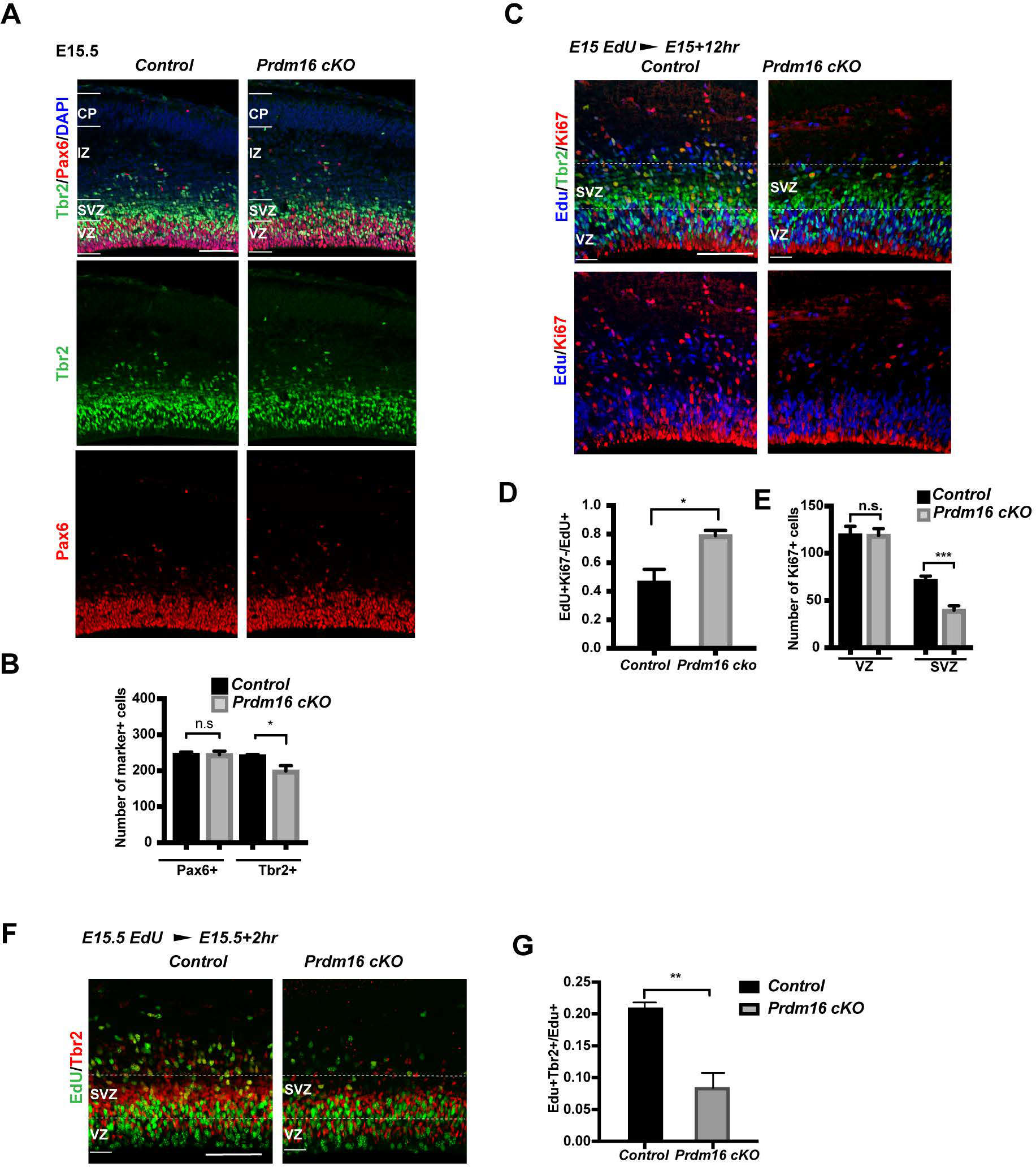
PRDM16 promotes IP cell proliferation during late cortical neurogenesis. **(A)** Images of E15.5 control and Prdm16 cKO cortices, stained with the IP marker Tbr2, the RG marker Pax6 and DAPI. **(B)** Quantification of the marker+ cells in 200 µm width column across the cortex (n=3). **(C)** Images of El5.5 control and Prdm16 cKO cortices, stained with EdU, Tbr2 and Ki67. White dashed lines highlight the SVZ. Scale bar: 50 µm. **(D)** Quantification of the fraction of EdU+Ki67-cells in 200 µm over EdU+ cells (n=3). **(E)** Quantification of Ki67+ cells in 300 µm (n=3). **(F)** Images of E15.5 with a 2-hour EdU pulse labeling, stained with EdU and Tbr2 antibodies. **(G)** Quantification of the fraction of EdU+Tbr2+ over EdU+ cells. Scale bar: 50 µm. All data are shown as mean+/−SD; *p<0.05; **p<0.01; ***p<0.001.

We next examined cell counts and proliferation of RG and IP cells at E13.5. In contrast to E15.5, neither the Pax6+ nor the Tbr2+ cells showed change in cKO cortex (**Supplemental Fig. S3A-B**). Staining with PH3 confirmed no change in the number of mitotic cells (**Supplemental Fig. S3C-D**). To test cell cycle exit rate, we injected pregnant mice with EdU at E13 and analyzed the cortex at E13.5. There was no significant change in the fraction of EdU+Ki67-cells over all EdU+ cells in cKO cortices (**Supplemental Fig. S3E-F**), indicating that cell exit did not occur earlier in the mutant at this stage. Another RG marker, Sox2, did not show any change (**Supplemental Fig. S3 F**).

Together, these findings suggest that PRDM16 regulates RG neurogenesis in a stage specific manner: first, it promotes the temporal transition of RG between E13.5 and E14.5; second, it promotes IP proliferation during late-neurogenesis.

### PRDM16 modulates levels of neuronal specification genes in RG

We hypothesized that PRDM16 may regulate the transcriptional program of RG. To this end, we generated RNA-seq data from E13.5 control and KO mutant forebrains (FB) (**Fig. 4A**). We identified 35 downregulated and 47 upregulated genes in KO versus control, using a cutoff of P value < 0.05 and fold-change > 1.5-fold (**Supplemental Fig. S4A-B**). We compared our FB RNA-seq data with the published RNA-seq data of sorted RG, IPs and cortical neurons (CNs) from E15.5 control and *Emx1^IREScre^*-mediated *Prdm16* cKO cortices (Baizabal et al. 2018). Most of the de-regulated genes in the mutant FB were also de-regulated in the E15.5 mutant RG (**Supplemental Fig. S4A**). Consistent with RG-specific expression of *Prdm16*, RG is the primary cell type where PRDM16 directly controls gene expression.

**Figure 4.**
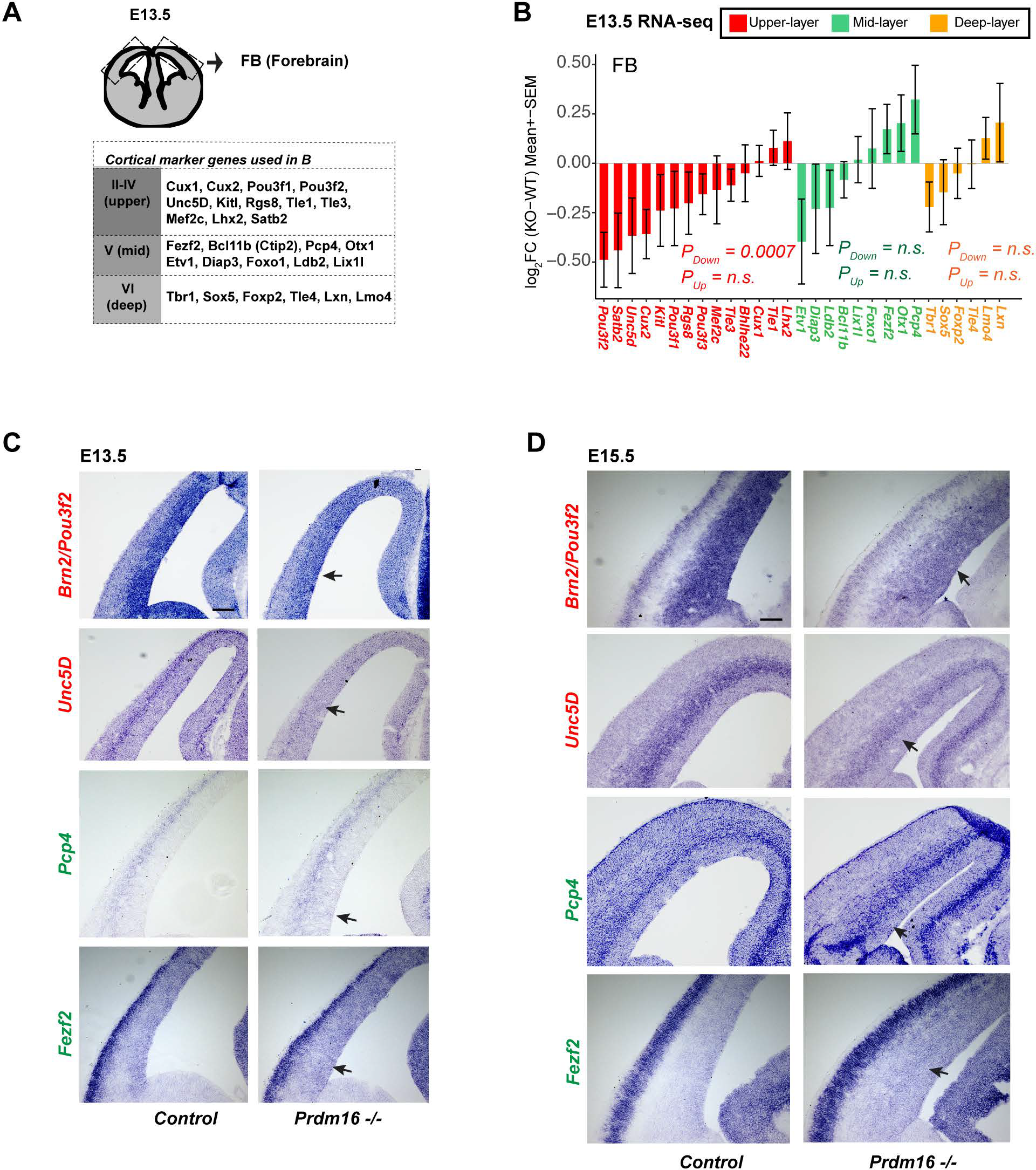
Depletion of Prdml6 led to defective gene expression in the developing cortex. **(A)** Schematic of the RNA-seq tissue and a list of layer marker genes described in the study. **(B)** Fold-changes of upper-, mid-, and deep-layer genes are shown. A gene set test shows the upper layer markers are significantly down-regulated. **(C-D)** In situ hybridization for Pou3f2/Brn2, Unc5D, Pcp4 and Fezf2 on control and mutant cortices at El 3.5 and El 5.5. Black arrows indicate Scale bar:100 µm.

We next examined expression of layer marker genes in the RNA-seq data (**Fig. 4A**, (Molyneaux et al. 2007; Zahr et al. 2018)). Several upper-layer genes showed decreased expression in the E13.5 KO FB (**Fig. 4B**). Using a limma-based gene set testing (See Methods), we confirmed significant down-regulation of the upper-layer markers as a gene set (*P*_down-regulation_ = 0.0007). Interestingly, given that the upper-layer neurons are not specified at E13.5, their expression changes likely occurred in progenitor cells.

Neither the mid-layer nor the deep-layer genes showed significant change as a group in the KO FB, despite the expression of three genes, *Pcp4*, *Otx1* and *Fezf2*, showing mild increase (**Fig. 4B**). We reasoned that at E13.5 the mid-and deep-layer neurons are specified and that many of the mid-layer genes are also expressed in the deep-layer, making it hard to reveal cell-type specific changes for these genes from whole FB data. To verify the changes, we selected *Pou3f2/Brn2 and Unc5D*, two upper-layer genes, and *Pcp4 and Fezf2*, two mid-layer markers for *in situ* hybridization experiments (**Fig. 4C-D**). Expression of *Brn2* and *Unc5D* was reduced in the mutant VZ and SVZ respectively, confirming the reduction of their expression in mutant progenitors. Expression of *Pcp4* and *Fezf2* showed an increase in the mutant VZ and SVZ, albeit the increase of *Fezf2* expression more obvious at E15.5. We also analyzed changes of the layer genes in the published E15.5 RG, IP and CN RNA-seq data (Baizabal et al. 2018) and observed a similar trend: the upper-layer genes are significantly downregulated in all three cell types (**Supplemental Fig. S4B**). The mid-layer genes as a group showed significant up-regulation in mutant RG and IPs, but not in mutant CN, suggesting these genes may be regulated in progenitors.

These results demonstrate that neuronal specification genes are already expressed in progenitors and that their normal expression is disrupted by *Prdm16* depletion.

### PRDM16 mainly functions as a transcriptional repressor in RG

To determine direct targets of PRDM16 in the developing brain and investigate how the targets are regulated, we performed chromatin immunoprecipitation followed by deep sequencing (ChIP-seq) from E13.5 heads. Using an IDR (Irreproducibility Discovery Rate) pipeline (see Methods), we identified 2319 confident peaks (IDR < 5%), of which 40% were mapped to intergenic regions, 30% to introns and only 20% close to promoters. This result indicates that PRDM16 mainly binds to distal enhancers (**Fig. 5A**). Gene Ontology analysis shows that PRDM16-bound genes are enriched for nervous system development, migration signaling and RG function (**Supplemental Fig. S5A-B**). We compared our E13.5 whole-head ChIP with the published E15.5 cortex ChIP data (Baizabal et al. 2018), and found around 30% (798) of the E13.5 peaks overlap with the E15.5 peaks (**Supplemental Fig. S5C**). The overlapping sites represent continuous binding by PRDM16 between E13.5 and E15.5.

**Figure 5.**
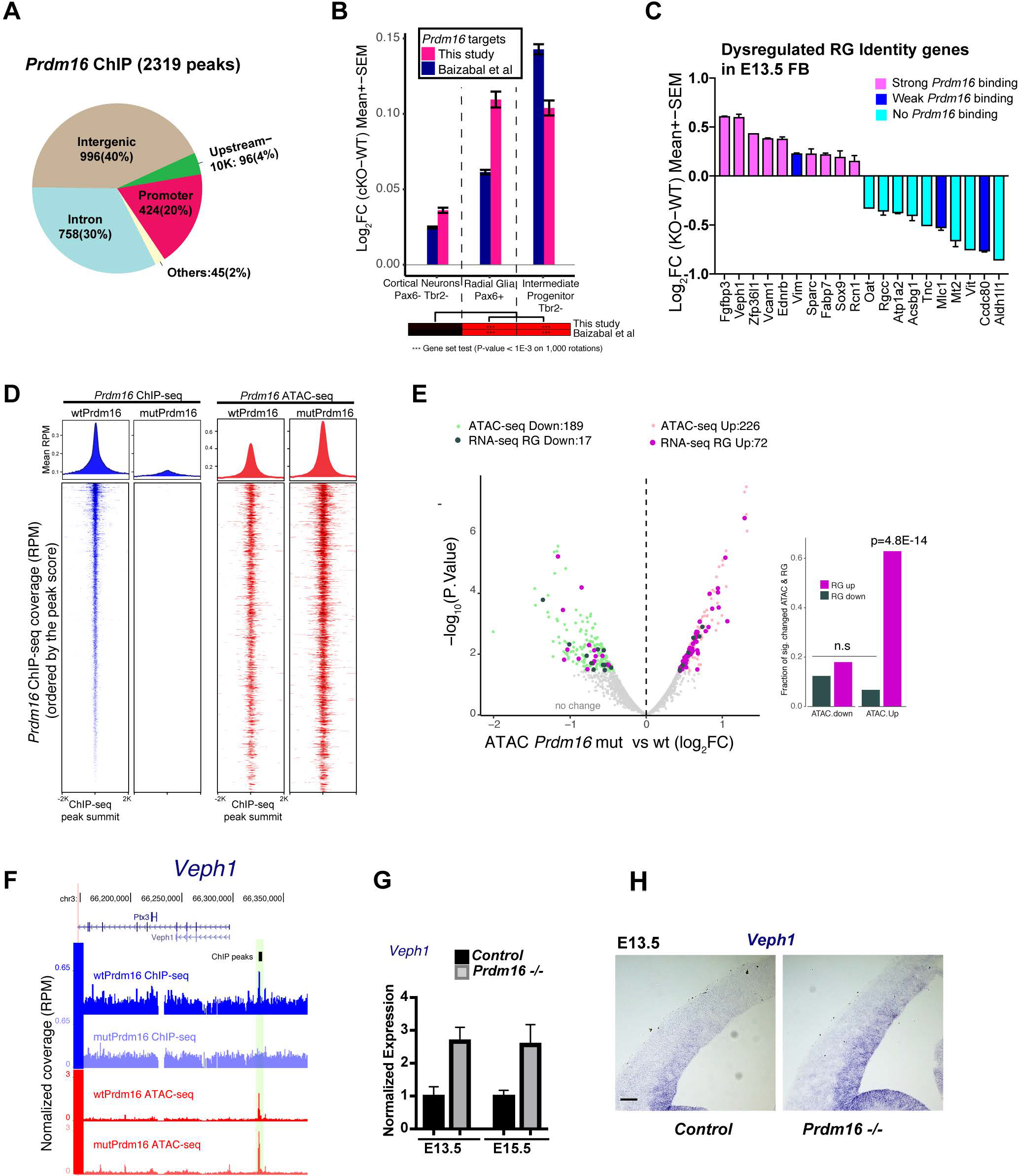
PRDM16 represses its target gene expression. **(A)** Genomic distribution of PRDM16 ChIP-seq peaks. **(B)** Gene set testing shows PRDM16 targets with a trend of up-regulation in cKO vs control in RG and IP, but not in CN. **(C)** A subset of the RG identity genes bound by PRDM16 show up-regulation in Prdm16 mutant FB while those weakly or not bound by Prdm16 were down-regulated. **(D)** The volcano plot shows significantly increased (light pink) or decreased (light green) ATAC-seq signal in mutant vs control at PRDM16-bound loci (FDR <= 0.2). The associated genes that had expression change in mutant RG were indicated in purple (up-regulated) or dark-green (down-regulated). The bar plot on the right side shows the fraction of genes that changed expression in mutant RG over all the genes that changed ATAC signals on the Prdm16-bound peaks. Gene up-regulation correlates with increased ATAC-seq signal. **(E)** Screenshot of the Veph1 gene locus, an example of bound and upregulated genes with increased chromatin accessibility**. (F-G)** RT-qPCR and in situ hybridization confirms de-repression of Veph1 in E13.5 and E15.5 KO FB.

We then analyzed how the targets are regulated by PRDM16. By applying gene set testing for all the targets as a set, we found that the targets in both E13.5 and E15.5 have a significant trend of de-repression in mutant RG (P <0.001) and IPs (P <0.001) but not in CNs (P=0.19) (**Fig. 5B**), suggesting that many targets are normally repressed by PRDM16. To confirm the finding from global analysis, we checked a group of genes called RG core identity genes (Yuzwa et al. 2017) highly expressed throughout neurogenesis. We found that 20 out of the 90 RG identity genes showed de-regulation in the *Prdm16* mutant FB. Only the subset bound by PRDM16 became upregulated in the mutant FB (**Fig. 5C**) or the mutant RG and IPs (**Supplemental Fig. S5D**).

We next examined chromatin states of E13.5 control and mutant cortices by using ATAC-seq (Assay for Transposase-Accessible Chromatin using sequencing) (Buenrostro et al. 2013) that measures chromatin accessibility. Higher ATAC-seq signals in the genome correlate with active cis-regulatory elements (Daugherty et al. 2017). At PRDM16-bound regions, there is high ATAC-seq intensity that became even higher in the mutant (**Fig. 5D**), suggesting that loss of *Prdm16* led to increased chromatin accessibility at its targeted sites. We quantified changes of ATAC-seq coverage on the PRDM16 ChIP-seq peaks between control and mutant. 226 and 189 peaks respectively showed increased and reduced coverage (FDR<0.2 and FC>1.4-fold) (**Fig. 5E, Supplemental Fig. S5E**). We then examined expression changes for the genes whose loci associate with accessibility changes. Interestingly, many of the up-regulated genes in E15.5 mutant RG (**Fig. 5E**) or mutant E13.5 FB (**Supplemental Fig. S5E**) had increased chromatin accessibility, whereas down-regulated ones do not show either trend. Validation on one of the genes, *Veph1*, by RT-qPCR and *in situ* hybridization confirmed de-repression of *Veph1* in *Prdm16* mutant (**Fig. 5F–H**). Hence, we conclude that PRDM16 primarily acts as a repressor in RG through maintaining accessibility of chromatin.

### PRDM16 directly represses mid-layer genes including Fezf2

Since PRDM16 represses transcription, as indicated above, expression of its targets in RG may be relatively low. We then asked in which cell types the target genes have higher expression. To address this, we first re-analyzed the published scRNA-seq data from E13.5 (Yuzwa et al. 2017) to obtain cell-type specific transcriptomes. We identified 6 clusters and assigned the cell type to each cluster (**Supplemental Fig. S6 A-B**) based on the presence of known markers. Consistent with the previous finding (Zahr et al. 2018), the RG and the IP clusters express many layer marker genes (**Supplemental Fig. S6B**). We then plotted the percentage of cells that contain high summed expression (normalized value > 180, see Method) of PRDM16 targets per cell in each cluster (**Fig. 6A and Supplemental Fig. S6C**). The mid- and deep-layer neuron clusters show the highest, suggesting many of the PRDM16 targets are highly expressed in mid-and deep-layer neurons.

**Figure 6.**
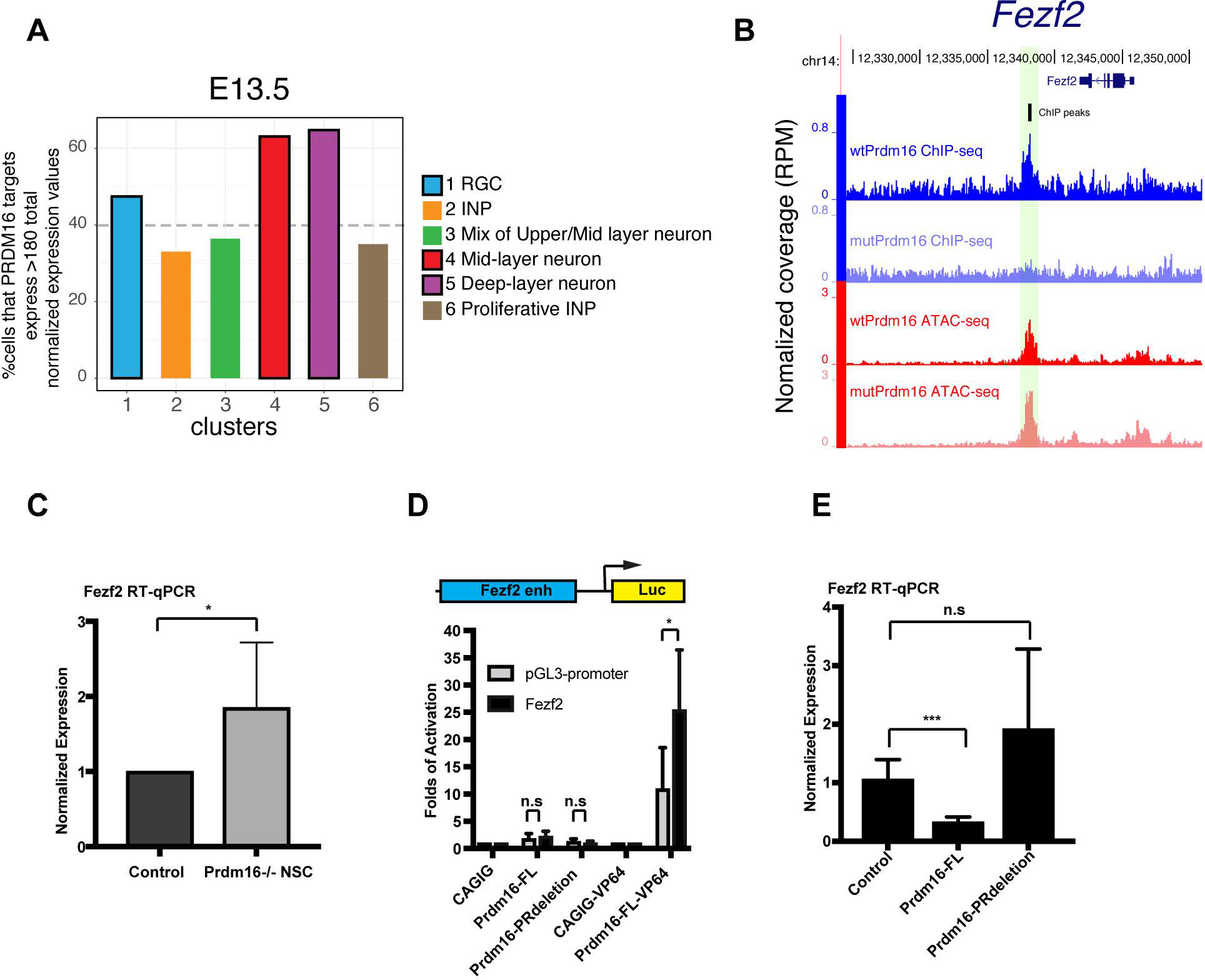
PRDM16 represses mid--layer neuronal genes including Fezf2. **(A)** Re-analysis of the cortex scRNA-seq data (Yuzwa et al. 2017) shows Prdml6 targets enriched in the RG, mid-and deep-layer neuron clusters at El 3.5. The Y-axis plots the percentage of cells that have summed expression of PRDM16 targets per cell (log2counts, normalized by library size, only the cells that have >180 expression value after normalization are included). **(B)** Screenshot of the Fezf2 locus with a PRDM16 peak in the RG enhancer of Fezf2. **(C)** RT-qPCRfrom primary neural stem cell culture of control and Prdml6 mutant cortical cells. Three pairs of control and Prdml6 KO embryos were used. **(D)** Luciferase assays in N2A cells. Prdml6-FL-VP64 significantly induced expression of the Fezf2 reporter but not the empty pGL3-luc alone. Four biological replicates were used. **(E)** RT-qPCRfrom the N2A cells expressing pCDH-Puro (empty vector control), Prdml6-FL or Prdml6-PRdeletion constructs. Two independent stable lines for each construct and three technique replicates of each stable line were used. All data are shown as mean+/−SD; *p<0.05; **p<0.01; ***p<0.001; n.s., not significant.

One of the PRDM16 targets is *Fezf2* (**Fig. 6B**), a mid-layer neuron determinant (Molyneaux et al. 2005). PRDM16 binds to the distal enhancers of *Fezf2*, one of which is known to drive *Fezf2* expression in RG (Shim et al. 2012). We confirmed *Fezf2* is de-repressed in *Prdm16* KO mutant neural stem cells, using primary NSC culture (**Fig. 6C**). We then tested responsiveness of this downstream RG enhancer to PRDM16 using a luciferase reporter driven by the *Fezf2* enhancer (**Fig. 6D**). PRDM16 fused with an VP64 activation domain induced higher expression of the *Fezf2* reporter compared to the pGL3-promoter alone, confirming PRDM16 binding to this enhancer. However, PRDM16 (PRDM16-FL) alone or the truncated version (PRDM16-PRdeletion, lack of the PR domain) did not have effect on the reporter (**Fig. 6D**). We reasoned that PRDM16 may require chromatin context for its regulatory activity which is lacking in transient transfection assay. To overcome this, we measured the endogenous level of *Fezf2* mRNA by RT-qPCR from N2A cells infected with a control construct or with either PRDM16-FL or PRDM16-PRdeletion (**Fig. 6E**). Endogenous *Fezf2* expression was reduced in cells expressing PRDM16-FL but not in those expressing PRDM16-PRdeletion, suggesting that PRDM16 needs endogenous chromatin context or other cis-regulatory element(s) to repress *Fezf2* and the PR domain is essential for its repressive activity.

### PRDM16 influences temporal dynamics of RG gene expression

We hypothesized that the gene expression program of E13.5 RG may differ from that of E15.5 RG and that PRDM16 may influence the dynamics of gene expression in RG. To test this, we first identified dynamic genes in RG, by performing differential expression analysis between the E13.5 and the E15.5 RG clusters from the published scRNA-seq data (see Method). 120 and 248 genes show higher expression at E13.5 and at E15.5 respectively (FDR < 0.2, FC>1.4-fold) (**Fig. 7A**). We then examined gene expression changes of these genes in *Prdm16* mutant RG (*Prdm16* cKO RNA-seq data (Baizabal et al. 2018)) and found 24 of them showing most significant changes (FDR < 0.05) (**Fig. 7B, Supplemental Fig. S7A**). All the up-regulated genes have PRDM16 binding, conforming the repressive activity of PRDM16. Among these genes, *Cdkn1c* encodes the cell cycle regulator p57^KIP2^ that suppresses progenitor cell proliferation in early neurogenesis (Mairet-Coello et al. 2012). Normal expression of *Cdkn1c* in RG decreases 2-fold from E13.5 to E15.5, and it is more strongly expressed in IP than in RG (**Supplemental Fig. S7B**), suggesting alleviation from its inhibitory activity in later stage may be required for higher proliferation of IPs. Interestingly, *Cdkn1c* was up-regulated in *Prdm16* mutant RG and IP at E15.5 but not in mutant FB at E13.5 (**Fig. 7C**), which may account for reduced proliferation of mutant IPs at E15.5. Another gene Flrt3 encodes fibronectin leucine rich transmembrane protein 3, a repulsive cue for the UNC5 family receptors in guiding cell migration (Yamagishi et al. 2011). Expression of Flrt3 increased in *Prdm16* mutant (**Fig. 7D**). As UNC5D is specifically expressed in upper-layer neurons, the potential action between UNC5D and FLRT3 provides a possible mechanism specific for upper-layer neuron migration. These results demonstrate that PRDM16 regulates expression of a subset of temporally-dynamic genes which may mediate its roles in promoting temporal transition of RG.

**Figure 7.**
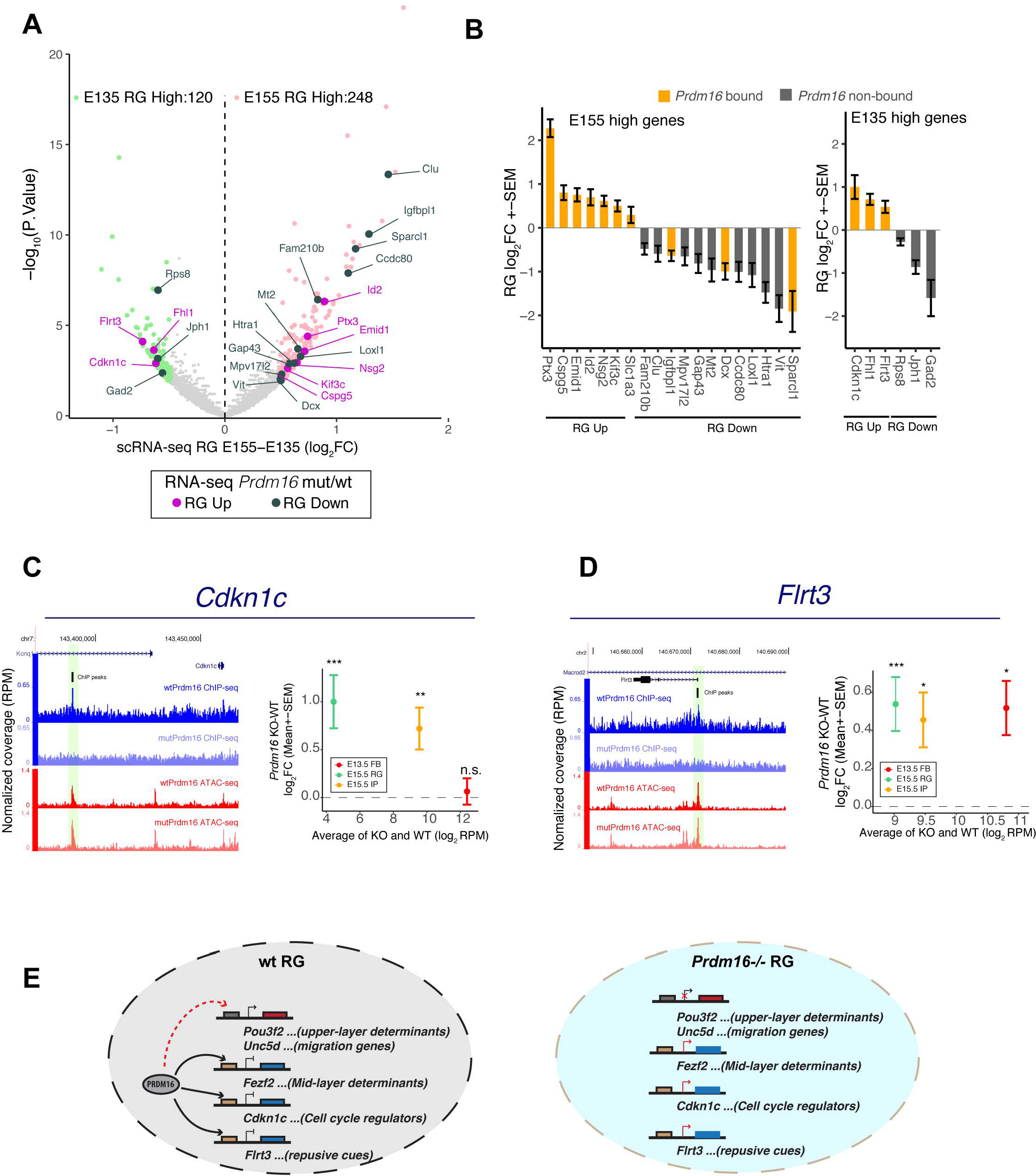
PRDM16 regulates temporal dynamics of RG gene expression. **(A)** Differentially expressed genes (FDR<0.2 and FC>1.4-fold) between E15.5 and E13.5 RG in the volcano plot include 248 increased (light pink) and 120 decreased (light green) genes in E15.5 versus E13.5 RG. The 24 most significantly up-and down-regulated genes were highlighted in purple and dark-green respectively in Prdm16 cKO/WT RG (FDR <0.05). **(B)** Expression changes of the 24 genes in Prdm16 cKO/WT RG were plotted. The genes containing PRDM16 binding peaks are highlighted in orange. **(C-D)** Screenshots and expression changes of two E13.5 high genes, Cdkn1c (C) and Flrt3 (D). Average RPM of KO and WT on x-axis shows absolute expression level. y-axis plots the fold-changes between KO/WT. *FDR<0.1; **FDR<0.05; *** FDR<0.01. **(E**) Proposed model of how PRDM16 controls RG neurogenesis through regulating different classes of genes. In Prdm16 mutant RG, de-repression of genes encoding mid-layer determinants, stage-specific cell-cycle regulators and migration cues consequently leads to prolonged production of mid-layer neurons, reduced IP proliferation and compromised neuronal production and migration for upper-layer neurons.

## Discussion

Our results demonstrate that PRDM16 is a critical transcriptional regulator that controls the gene expression program of RG during cortical neurogenesis. Regulation by PRDM16 is required for the timed progression of RG between early and late phases of neurogenesis. PRDM16 executes the temporal shift by establishing the stage-specific gene expression program including neuronal specification genes, cell cycle regulators and genes for upper-layer neuron migration.

Recently Baizabal *et al* reported that PRDM16 regulates upper-layer neuron production and migration but does not affect deep-or mid-layer neuron fate (Baizabal et al. 2018). However, we found that there is prolonged production of mid-layer neurons in *Prdm16* cKO cortex in addition to its effects on upper-layer neurons. The discrepancy may result from the methods used to assess cell fates. We distinguished L5 mid-layer neurons from L6 deep-layer neurons while they assessed L5 and L6 neurons as a whole population. We dissected the regulatory network controlled by PRDM16 at the transition of mid-to late-neurogenesis and identified different classes of genes responsible for PRDM16 functions. We propose a model that includes three key points (**Fig. 7E**): 1) PRDM16 represses mid-layer determinants to allow timely upregulation of upper-layer genes in RG; 2) PRDM16 represses cell cycle inhibitors to allow higher proliferation of IPs at later neurogenesis; 3) PRDM16 controls genes encoding guidance cues needed for upper-layer neuronal migration.

### PRDM16 and the temporal identity of RG

The understanding of temporal control of RG has been augmented over the years. A number of transcription factors and epigenetic regulators have been shown to control the timing of cortical neuronal specification (McConnell and Kaznowski 1991; Chenn and Walsh 2002; Fukumitsu et al. 2006; Ge et al. 2006; Shen et al. 2006; Hsu et al. 2015; Dennis et al. 2017). Progressive hyperpolarization of the membrane of RG regulates the sequential generation of neuronal subtypes through modulating Wnt signalling (Vitali et al. 2018). Moreover, it was recently revealed that RG are primed with a spectrum of neuronal genes. Post-transcriptional mechanisms, including translational repression (Zahr et al. 2018) and the N6-methyladenosine (m6A) RNA modification (Yoon et al. 2017), regulate RG progression through preventing precocious production of neuronal proteins in RG. Some questions still remain. For example, how is the priming status of RG established? How does the pre-established transcriptional program impact daughter cell fate? We found that *Prdm16* mutant RG show disrupted expression of mid-and upper-layer genes and several temporally-dynamic genes involved in proliferation (*Id2, Cdkn1c*) and migration (*Flrt3, Dcx, Sparcl1*). These results suggest that PRDM16 may be involved in setting up the primed gene expression program of RG. PRDM16 is expressed throughout cortical neurogenesis. A question is how its activity is triggered at the onset of mid-to late-neurogenesis transition. Interestingly, *Prdm16* co-clusters with *Slc1a3*, a regulator of metabolism of glutamate and ion flux(Vandenberg and Ryan 2013). It will be of interest to test a potential function of SLC1A3 in integrating extrinsic and intrinsic signals.

Notably, Hamlet, the ortholog of PRDM16 controls the temporal identity of intermediate progenitors in *Drosophila* neuroblast lineage (Eroglu et al. 2014), suggesting that the role of the PRDM16 proteins are evolutionarily conserved.

### Transcriptional activity of PRDM16 in cortical development

PRDM16 can act as a repressor or an activator depending on its associated partners (Chi and Cohen 2016). We showed that PRDM16-bound genes have a trend of de-repression in *Prdm16* mutant, indicating its repressive role in the neocortex. The fact that many of the PRDM16 targets are expressed in RG suggest that repression by PRDM16 is not to fully silence genes but to maintain gene expression at the right level. In support of this, PRDM16 binding associates with open chromatin. Moreover, we do not rule out the possibility of PRDM16 being an activator, as our ChIP and RNA-seq data identified a small subset of genes that show PRDM16 binding and down-regulation in mutant.

The PR domain of PRDM16 is essential in repressing *Fezf2*. Baizabal *et al* showed that PRDM16 without the PR domain failed to rescue target gene de-repression (Baizabal et al. 2018). The PR domain of PRDM16 was shown to be essential in suppressing MLLr1 leukemia via intrinsic H3K4 methylation activity (Zhou et al. 2016). We did not observe global changes of H3K4me1 or H3K4me2 levels in mutant cortex by immunostaining (data not shown). In agreement with this result, Baizabal *et al* did not detect significant change of H3K4 methylation levels using ChIP-seq (Baizabal et al. 2018). Future studies are needed to address the mechanistic nature of how the PR domain or any other domain contributes to the function of PRDM16 in the neocortex.

*Prdm16* is among the many genes deleted in human 1p36 microdeletion syndrome, a disorder that displays a wide variety of disease conditions. According to the previous identified function of PRDM16 in normal development, loss of *Prdm16* might contribute to several problems including the craniofacial and cardiac defects and hydrocephalus of the syndrome (Bjork et al. 2010; Arndt et al. 2013; Shimada et al. 2017). Our findings, along with the study by Baizabel *et al*, defined a mechanism by which *Prdm16* loss of function in the formation of Heterotopia, a neurodevelopmental disorder that leads to severe mental retardation and seizures that were also seen in the 1p36 syndrome. More mechanistic insights of PRDM16 function will increase our understanding of its developmental roles in cell fate specification and its pathological role in diseases.

## Materials and methods

### Animals and processing

All animal procedures were approved by Swedish agriculture board (Jordbruks Verket) with document number Dnr 11553-2017 and the MWU Institutional Animal Care and Use Committee. The *Prdm16*^cGT^ and *Prdm16^cGTreinv^* mice (Strassman et al. 2017) were maintained by outcrossing with the FVB/NJ line. *B6.129S2-Emx1tm1(cre)Krj/*J (*Emx1^IREScre^*) (Gorski et al. 2002) were used to generate conditional gene trap knockout animals as described previously (Strassman et al. 2017).

### Molecular cloning

The pCAGIG plasmid (Addgene) was inserted with a fragment encoding a nuclear localization signal (NLS) and 3xFlag in the EcoRI site. To make pCAGIG-Prdm16-FL or pCAGIG-Prdm16-PRdeletion, the full-length open reading frame (ORF) or the truncation that lacks coding sequence for amino acid 2 to amino acid 180 of *Prdm16* was PCR amplified from MSCV-Prdm16 (Addgene 15504) and inserted between the EcoRI and XhoI sites in pCAGIG-NLS-Flag. The VP64 fragment was then inserted to the XhoI site of pCAGIG-Prdm16 to make pCAGIG-FL-VP64.

The plasmids used for making stable cell lines, pCDH-Prdm16 and pCDH-Prdm16-PRdeletion, were generated as follow: The Prdm16 FL ORF, the PR-deletion coding sequences or the NLS-3xFlag was digested from their pCAGIG plasmids and inserted sequentially to the pCDH-CMV-MCS-EF1-Puro plasmid (System Biosciences) between the EcoRI and NotI sites (for Prdm16-FL and Prdm16-PRdeletion) and the XbaI and EcoRI sites (for NLS-3xFLAG).

### Immunochemistry, BrdU and EdU labelling and confocal imaging

At designed stages, embryos or pulps were perfused with PBS followed by 4% paraformaldehyde. The perfused brains were dissected, fixed overnight and sectioned coronally using a vibratome (Leica Microsystems). Immunostaining was done according to standard protocols as previously used (Dai et al. 2013a). The list of primary secondary antibodies and using condition is provided in supplemental table 4.

BrdU (5-bromo-2'-deoxyuridine) and EdU (5-ethynyl-2′-deoxyuridine) (5-20 µg/g of body weight) were injected into the peritoneal cavity of pregnant mice. BrdU incorporation was measured by immunostaing using an antibody against rat-BrdU (Abcam) and mouse-BrdU (DSHB). EdU incorporation was detected with the Click-iT assay (Invitrogen) according to the manufacturer’s instructions. Imaging was done on Zeiss confocal microscope. ZEN (ZeissLSM800), ImageJ (NIH) and Photoshop (Adobe) were used for analysis and quantification.

### In situ hybridization

The mouse brains at defined ages were dissected and fixed for 12 hours in 4% PFA, cryoprotected in 25% sucrose overnight, embedded in O.C.T, and sectioned at 18 µm on Leica cryostatsCM3050s. RNA *in situ* hybridization was performed using digoxigenin-labeled riboprobes as described previously. Detailed protocols are available upon request. Images were taken using a Leica DMLB microscope.

### Quantification and statistical analysis

Cell numbers were manually counted in ImageJ/Fiji cell counter (National Institute of Health, USA). Number of marker positive cells in the control and KO mutant at P0 were determined by counting the average number of positive cells in three 80 *µm* width columns. Number of marker positive cells in the control and KO or cKO mutants at E15.5 were determined by counting number of positive cells in 100 *µm* width column from layer IV to VI. Numbers of cells in the control and cKO cortex at P15 were determined by counting the number of positive cells in one 250um width column within the whole cortex in two different areas (medial and dorsal lateral). For proliferation analysis at E15.5, numbers of Pax6+, Tbr2+, EdU+, Ki67+ cells were determined by counting the total number of positive cells in two 100 *µm* width columns in both VZ and SVZ. For proliferation analysis at E13.5, numbers of Pax6+, Tbr2+, EdU+, Ki67+ cells were determined by counting the total number in 300 *µm* width cortex. The production of daughter neurons was reflected by the cell cycle exit through measuring the ratio of EdU+Ki67-cells in total EdU+ cells. Numbers of BrdU+Ctip2+ or BrdU-Ctip2+ cells at P5 were determined by counting number of positive cells in 300 *µm* width column in layer II to layer V. All data are presented as mean ± SD, and statistical significance was determined using two-tailed unpaired Student’s t test.

### Neural stem cell culture and RT-qPCR

Control and mutant embryonic cortices were dissected and dissociated into single cell suspension and digested with Acutase (Sigma). Cells were maintained in proliferation media (STEMCELL Technologies). 3 control or 3 *Prdm16* mutant neural stem cell cultures were grown for two days before RNA extraction by use of TRIzol reagent (Invitrogen). 4 ug of total RNA was further cleaned with Turbo DNase (Ambion) and used in reverse-transcription with RT master mix (ThermoFisher). To ensure the absence of genomic DNA, control qPCR was performed on a mock-reverse-transcribed RNA sample. Primer sequences are listed in Supplemental Table 4.

### Cell culture and Luciferase assays

The neuroblastoma cell line Neuro-2A (N2A) cells were cultured in 50% of DMEM (GIBCO) containing 10% fetal calf serum and 50% of optimen serum reduced medium. For luciferase assays, transfections were performed in 96-well plate using FugeneHD tranfection reagent (Promega). The following DNA combinations were used: 20ng of *Fezf2* luciferase reporter or the pGL3 promoter vector, 100ng of pCAGIG-Prdm16, pCAGIG-Prdm16PRdeletion, pCAGIG-Prdm16-VP64, pCAGIG-VP64 or pCAGIG. 2ng of Renilla-luciferase construct was used as internal control. After 24-hour incubation, transfected cells were lysed and luciferase activity was measured using Dual Luciferase Assays (Promega), and promoter activity was defined as the ratio between the firefly and *Renilla* luciferase activities.

For generating the cell lines that stably express control, Prdm16-FL or Prdm16-PRdeletion, lentiviral particles were first produced in 293T cells and then added to N2A cells for infection. The cells that stably express the corresponding constructs were selected and maintained in medium that contains puromycin. Two individual stable lines were generated for each of the constructs used in RT-qPCR analysis of the Fezf2 gene.

### ChIP-seq analysis

In each replicate, three E13.5 control or Prdm16 KO mutant heads were pooled, fixed and lysed. ChIP was performed as previously described (Dai et al. 2013b). DNA libraries were made using the NEBNext Ultra™ II DNA Library Prep Kit and sequenced on the Illumina Hiseq2500 platform.

The replicated *Prdm16* KO (x3) and control (x3) ChIP-seq samples, after the adaptor trimming by Trimmomatic, were mapped to the UCSC *Mus musculus* (mm10) genome assembly using Bowtie2 with the default parameters. The uniquely mapped reads (with mapping quality >= 20) were used for further analysis. The PRDM16 peaks were called by HOMER (v4.10) (Heinz et al. 2010). The peak replicate reproducibility was estimated by Irreproducibility Discovery Rate (IDR), using the HOMER IDR pipeline (https://github.com/karmel/homer-idr). As suggested by the Encode IDR guideline that IDR requires to initially call peaks permissively for the replicates, we used a relatively relaxed parameter “-F 2-fdr 0.3 -P .1 -L 3 -LP .1” for the true/pseudo/pooled replicates by the HOMER peak calling. The final confident peaks were determined by an IDR < 5%. The peaks that were overlapped with mm10 blacklist were also removed. For comparisons, we re-analyzed the *Prdm16* control and cKO ChIP-seq public data ((Baizabal et al. 2018); GSE111657) using the same HOMER IDR pipeline.

### RNA-seq differential expression analysis

Cortices of control and Prdm16 KO mutant E13.5 embryos were dissected for RNA extraction using Trizol reagent (Invitrogen). RNA quality of three biological replicates was tested by Agilent Bioanalyzer. RNA-seq libraries were made using the Illumina Truseq Total RNA library Prep Kit LT. Sequencing was performed on the Illumina Hiseq2500 platform.

After trimming the adaptor sequences using Trimmomatic, we mapped RNA-seq reads from the replicated *Prdm16* wild type (x3) and mutant samples (x3) to the UCSC *Mus musculus* (mm10) genome assembly using HISAT2. We normalized RNA-seq by the “Relative Log Expression” method implemented in the DESeq2 Bioconductor library (Love et al. 2014). Gene annotation was obtained from the iGenomes UCSC *Mus musculus* gene annotation. Differentially expressed mRNAs between *Prdm16* mutants versus wild type were identified, and FDR (Benjamini-Hochberg) was estimated, using DESeq2. For comparisons, we re-analyzed the differential expression of *Prdm16* WT and cKO RNA-seq public data (Baizabal et al. 2018; GSE111660) using the same method as above. The genes with P-value ≤ 0.05 were considered to be differentially expressed.

### ATAC-seq analysis

The ATAC-seq libraries were made according to the published method (Buenrostro et al. 2013) and using the Illumina Nextera DNA library kit. In brief, cortices were dissected from 3 control and 3 *Prdm16* KO E13.5 brains. Tn5 enzyme reaction was performed at 37 degrees for 30mins, followed by DNA purification. 11 cycles of PCR amplification was performed using barcoded adaptors and primers on purified DNA template. Libraries were purified and pooled before sequencing with illumina Next-seq platform. The replicated *Prdm16* KO (x3) and control (x3) ATAC-seq samples, after the adaptor trimming by Trimmomatic, were mapped to the UCSC *Mus musculus* (mm10) genome assembly using Bowtie2 with the default parameters. The high quality and uniquely mapped reads (with mapping quality >= 20) were used for further analysis. ATAC-seq differential expression analysis between Prdm16 mutants and wild types on the Prdm16 bound ChIP-seq peaks were performed by Limma R package.

The ATAC-seq peak calling was performed by HOMER using the “broad peak” option with parameters “-region -size 1000 -minDist 2500”, separately for the mutant and wild type. To compare active enhancers between E13.5 and E15.5, we further re-analyzed the publicly available histone mark H3K27ac Prdm16 ChIP-seq data at E15.5. We called the peaks against Input using “narrow peak” option by HOMER with the default parameters.

### Gene set enrichment testing

To test whether a set of genes are significantly changed amongst the differentially expressed (DE) genes from *Prdm16* wild type and mutant RNA-seq data, we used gene set testing function “camera” and “mroast” in the R limma package (Ritchie et al. 2015). We used “camera”, a ranking based gene set test accounting for inter-gene correlation, to test whether the layer markers are significantly changed as a set. We used “mroast” (number of rotations = 1000), a self-contained gene set test, to test whether the majority of the genes amongst PRDM16 targets are significantly up-or down-regulated. We also used “mroast to test which Gene Ontology (GO) terms and Reactome pathways are significantly up-or down-regulated in *Prdm16* mutant versus wild type.

### scRNA-seq analysis

To gain insights into cell types of the *Prdm16* targets, we reanalyzed the murine cortical time-series scRNA-seq data (Yuzwa et al. 2017). We employed the Bioconductor scRNA-seq analysis workflow for droplet-based protocols (Lun et al. 2016). (i) The cortical cells (the cells expressing Emx) were selected for the analysis. The low quality cells were first removed if they are 3 MAD (the median absolute deviation) lower than the median library size OR if they are 3 MAD lower the median gene expression OR if they are 4 MAD higher than the median mitochondrial reads. (ii) We used the deconvolution approach, a method to handle high zeros in scRNA-seq, to compute size factors for cells for normalization. (iii) The cells were constructed into graphs by constructing a shared nearest neighbor graph (SNN) and clustered by Walktrap algorithm. (iv) We manually assigned cell types to the identified clusters in each stage using the known neuron and layer markers.

To identify differentially expressed genes between E13.5 and E15.5, RG cells were extracted from E13.5 and E15.5 RG clusters and differential expression analysis was performed using edgeRQLF R package. The genes with FDR <= 0.2 and FC > 1.4-fold were considered to be significantly differentially expressed between E13.5 and E15.5.

## Supporting information

Supplemental Figures and list

## Acknowledgements

We thank the animal experimental core facility (ECF) and the imaging facility (IFSU) of Stockholm University and the National Genomic Institute of Scilife Laboratories, Sweden, for providing service and support. W.H was supported by the visiting scientist fellowship from China Scholarship Council. J.W. was supported by the Australian Research Council (ARC) Future Fellowship (FT60100143). The project was supported by the Young Investigator grant from Swedish Research Council (Vetenskapsrådet, 2014-5584) and the research grant from Swedish Cancer funding agency (CAN 2017/529) to Q.D.

## Author contributions

Q.D. conceived and designed the project. L.H. performed most of the experiments, with help from J.J, B.B and W.H. J.W performed all the computational analysis. Q.D., J.W. and L.H. analysed and interpreted the data. Q.D. wrote the manuscript with input from the other authors.

