## Supplemental Figures and list for "PRDM16 establishes lineage-specific transcriptional program to promote temporal progression of neural progenitors in the mouse neocortex"

**Supplemental materials**

**Figure S1:** Basic information of the PRDM16 protein and the Prdm16 gene-trap line for generating null or conditional knockout animals: relevant to Figure 1

**Figure S2:** *Prdm16* forebrain specific knockout animals display neuronal migration defect: relevant to Figure 2

**Figure S3:** PRDM16 regulates RG proliferation at E13.5: relevant to Figure 3

**Figure S4:** Top de-regulated genes in the mutant forebrains from RNA-seq analysis: relevant to Figure 4

**Figure S5:** Additionaly analysis on genes that associate with PRDM16 peaks: relevant to Figure 5

**Figure S6:** Re-analysis of single cell expression data: relevant to Figure 6

**Figure S7:** Differently expressed genes between E15.5 and E13.5 RG cells: relevant to Figure 7

**Supplementary Table 1**. All genes in FB RNA-seq data.

**Supplementary Table 2**. All genes from PRDM16 ChIP-seq calls using E13.5 whole head ChIP-seq (this study) and E15.5 cortex ((Baizabal et al. 2018))

**Supplementary Table 3**. Differentially expressed genes between the E13.5 and E15.5 RG clusters (re-analyzed from (Yuzwa et al. 2017))

**Supplementary Table 4**. Reagents and oligonucleotide sequences.

Baizabal JM, Mistry M, Garcia MT, Gomez N, Olukoya O, Tran D, Johnson MB, Walsh CA, Harwell CC. 2018. The Epigenetic State of PRDM16-Regulated Enhancers in Radial Glia Controls Cortical Neuron Position. *Neuron* **98**: 945-962 e948.

Yuzwa SA, Borrett MJ, Innes BT, Voronova A, Ketela T, Kaplan DR, Bader GD, Miller FD. 2017. Developmental Emergence of Adult Neural Stem Cells as Revealed by Single-Cell Transcriptional Profiling. *Cell Rep* **21**: 3970-3986.

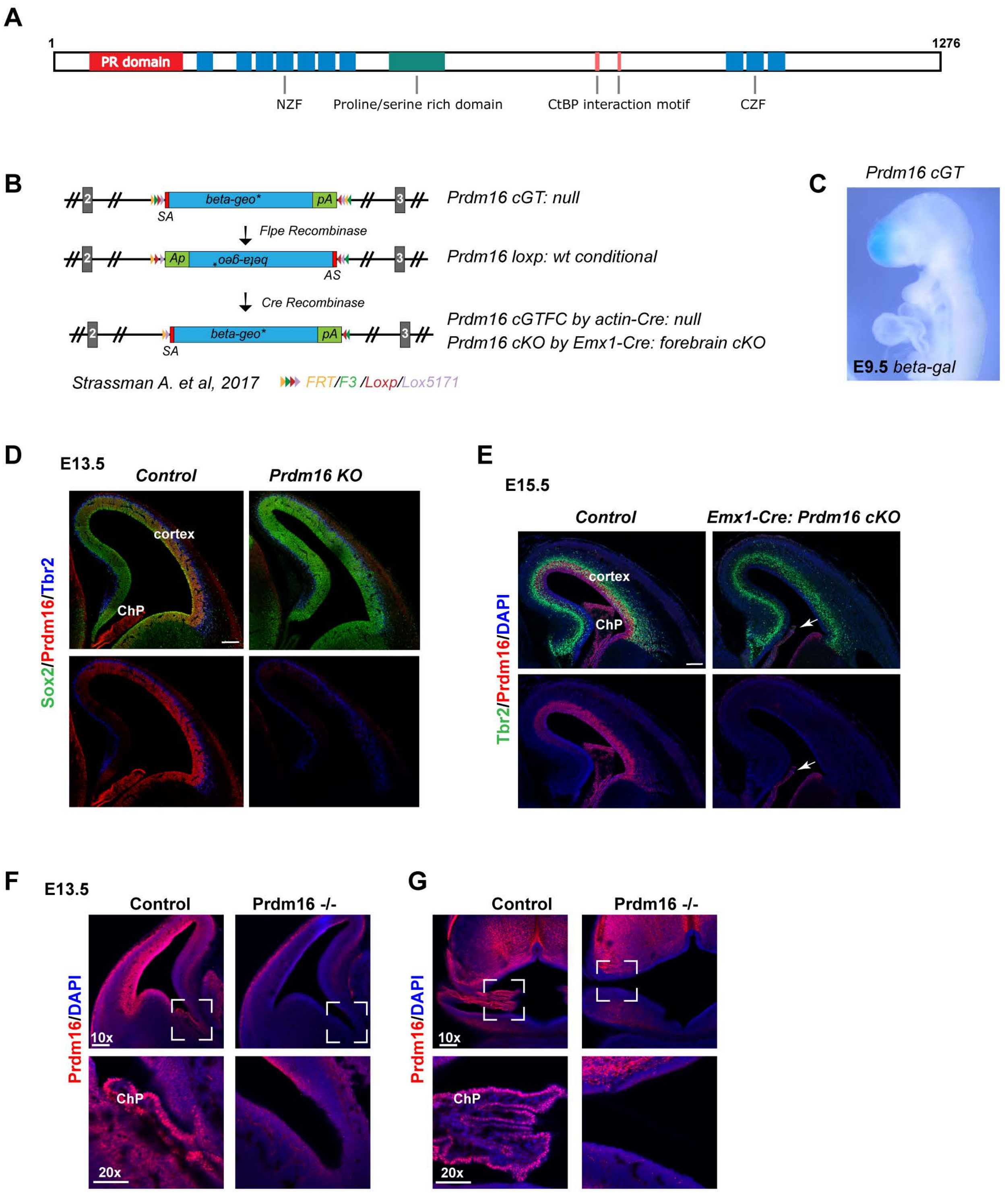
A
1
1276
PR domain
NZF
Proline/serine rich domain
CtBP interaction motif
CZF
B
beta-geo*
pA
SA
Prdm16 cGT: null
Flpe Recombinase
Ap
beta-geo*
AS
Prdm16 loxp: wt conditional
Cre Recombinase
beta-geo*
pA
SA
Prdm16 cGTFC by actin-Cre: null
Prdm16 cKO by Emx1-Cre: forebrain cKO
Strassman A. et al, 2017
FRT/F3 /Loxp/Lox5171
C
Prdm16 cGT
E9.5 beta-gal
D
E13.5
Control
Prdm16 KO
cortex
ChP
Sox2/Prdm16/Tbr2
E
E15.5
Control
Emx1-Cre: Prdm16 cKO
cortex
ChP
Tbr2/Prdm16/DAPI
F
E13.5
Control
Prdm16 -/-
Prdm16/DAPI
10x
ChP
20x
G
Control
Prdm16 -/-
Prdm16/DAPI
10x
ChP
20x

**Figure S1: Basic information of the PRDM16 protein and the *Prdm16* gene-trap line for generating null or conditional knockout animals and loss of the ChP in *Prdm16* KO mutant**

**(A)** Schematic of the largest isoform of the Prdm16 protein shows different functional domains of Prdm16. **(B)** Schematics of the gene-trap lines. **(C)** X-gal staining of a E9.5 embryo with the gene-trap ($Prdm16^{cGT}$) indicates that Prdm16 is expressed at E9.5. **(D)** Coronal section slices of control or $Prdm16^{cGT}$ KO mutant, stained with PRDM16 in red, the RG marker Sox2 in green and the IP marker Tbr2 in blue. PRDM16 staining is completely lost in the entire brain. Scale bar: 100 $\mu m$. **(E)** Coronal section slices of control or $Emx1^{IREScre}$-mediated *Prdm16* conditional mutant (*Prdm16* cKO), stained with Prdm16 in red, Tbr2 in green and the nuclei marker DAPI in blue. Prdm16 staining is mostly lost in the forebrain but remained in the ventral telencephalon and the ChP. White arrows indicate the presence of the ChP in the muant. **(F, G)** Images of E13.5 cortices stained with the Prdm16 antibody and DAPI, showing high expression of Prdm16 in the ChP in the control but absent in the Prdm16 null mutant in the lateral ventricle **(F)**, and in the 3rd ventricle **(G).** Lower panels are higher magnification images of the white square area of the upper panels. Scale bar: 50 $\mu m$.

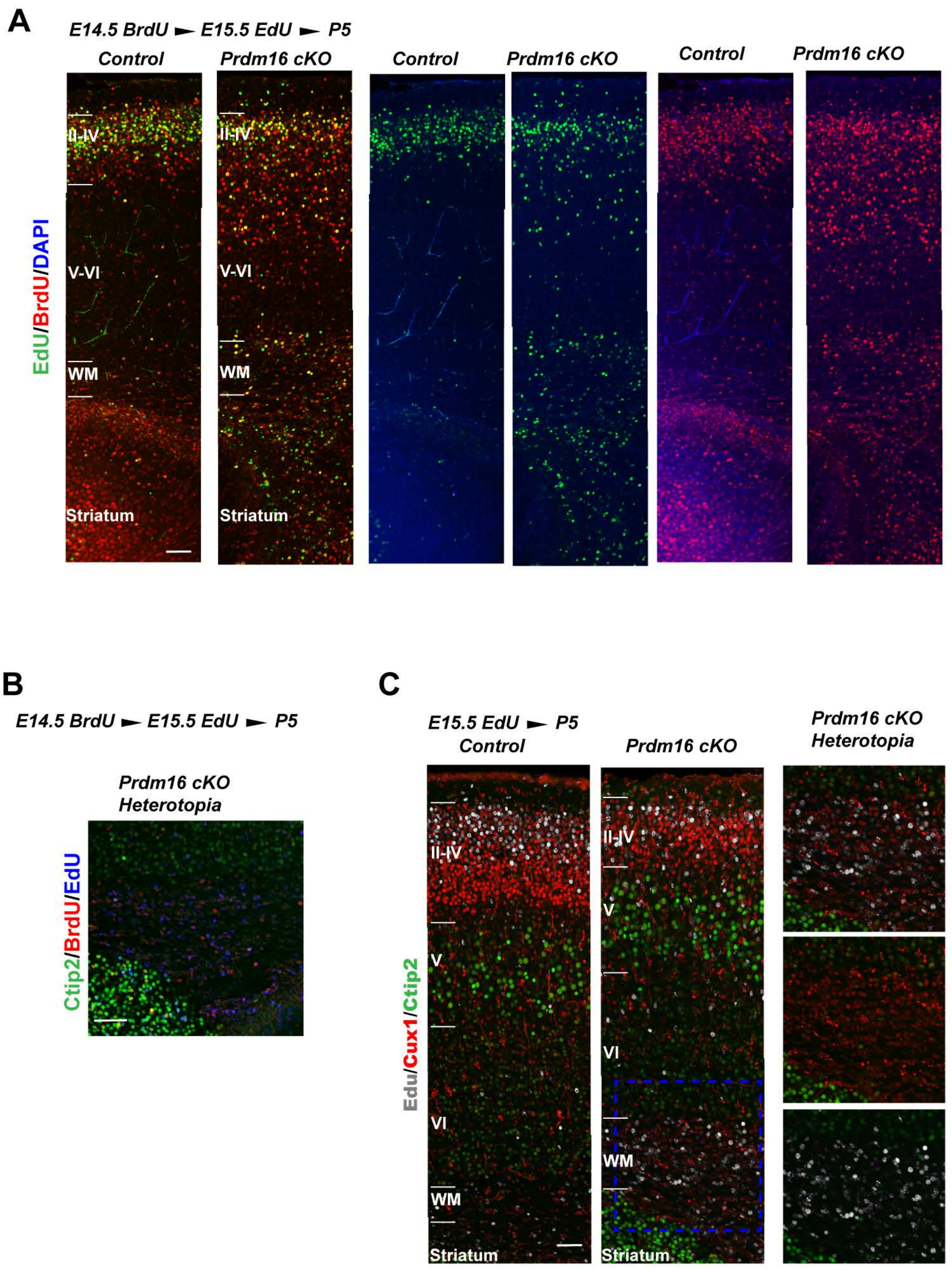


**Figure S2: *PRDM16* forebrain specific knockout animals display neuronal migration defect.** **(A)** Coronal sections of P5 control and cKO brains, stained with Brdu in red, Edu in green and the nuclei marker DAPI in blue. The cells labeled with Brdu or Edu were retained in the deep layer and in the heterotopia tissue in mutant cortex. **(B)** The heterotopia area in the cKO section, stained with Ctip2 in green, Brdu in red and Edu in blue, showing lack of Ctip2+ and Brdu+ or Edu+ double positive cells in heterotopia. **(C)** Coronal sections of control and cKO brains, stained with Cux1 in red, Edu in gray and Ctip2 in green. All of Edu+ cells are positive for Cux1. **B-C** together shows failure of migration is in the neurons co-stained with upper-layer marker Cux1 but not with mid-layer marker Ctip2.

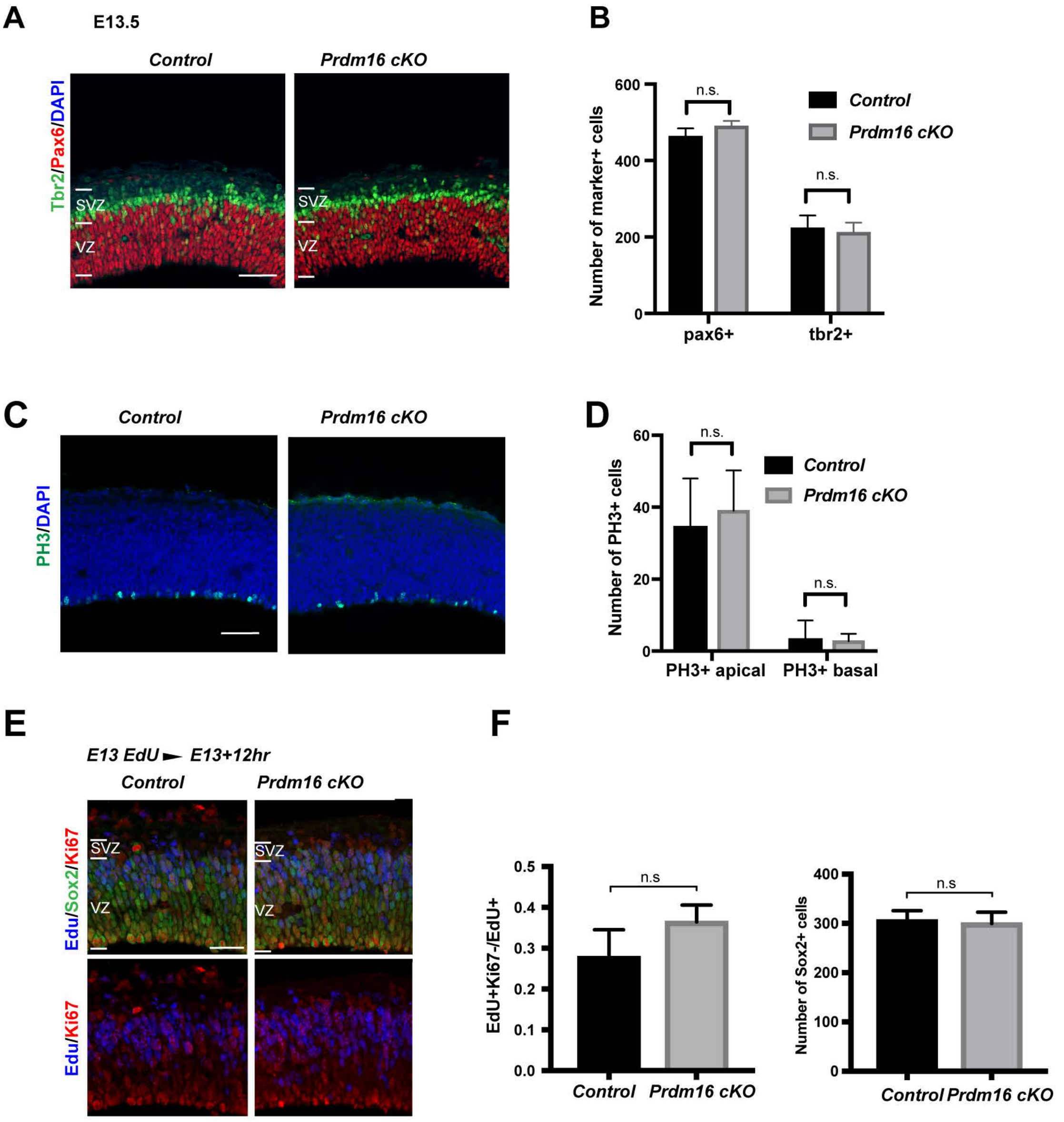


**Figure S3: PRDM16 regulates RG proliferation at E13.5**

**(A)** Images of E13.5 control and cKO brains, stained with Pax6 in red, Tbr2 in green and DAPI in blue. Scale bar: 50 $\mu m$. **(B)** Quantification of Pax6+ or Tbr2+ cells across the entire VZ and SVZ. **(C)** Images of E13.5 control and cKO brains, stained with the mitotic marker PH3. Scale bar: 50 $\mu m$. **(D)** Quantification of PH3+ cells across the entire VZ and SVZ. **(E)** Cell cycle exit analysis between E13 and E13.5. Scale bar: 50 $\mu m$. **(F)** Quantification of the fraction of Ki67-EdU+ relative to EdU+ cells (n=4). All data are shown as mean +/- SD; *p<0.05; **p<0.01; ***p<0.001.

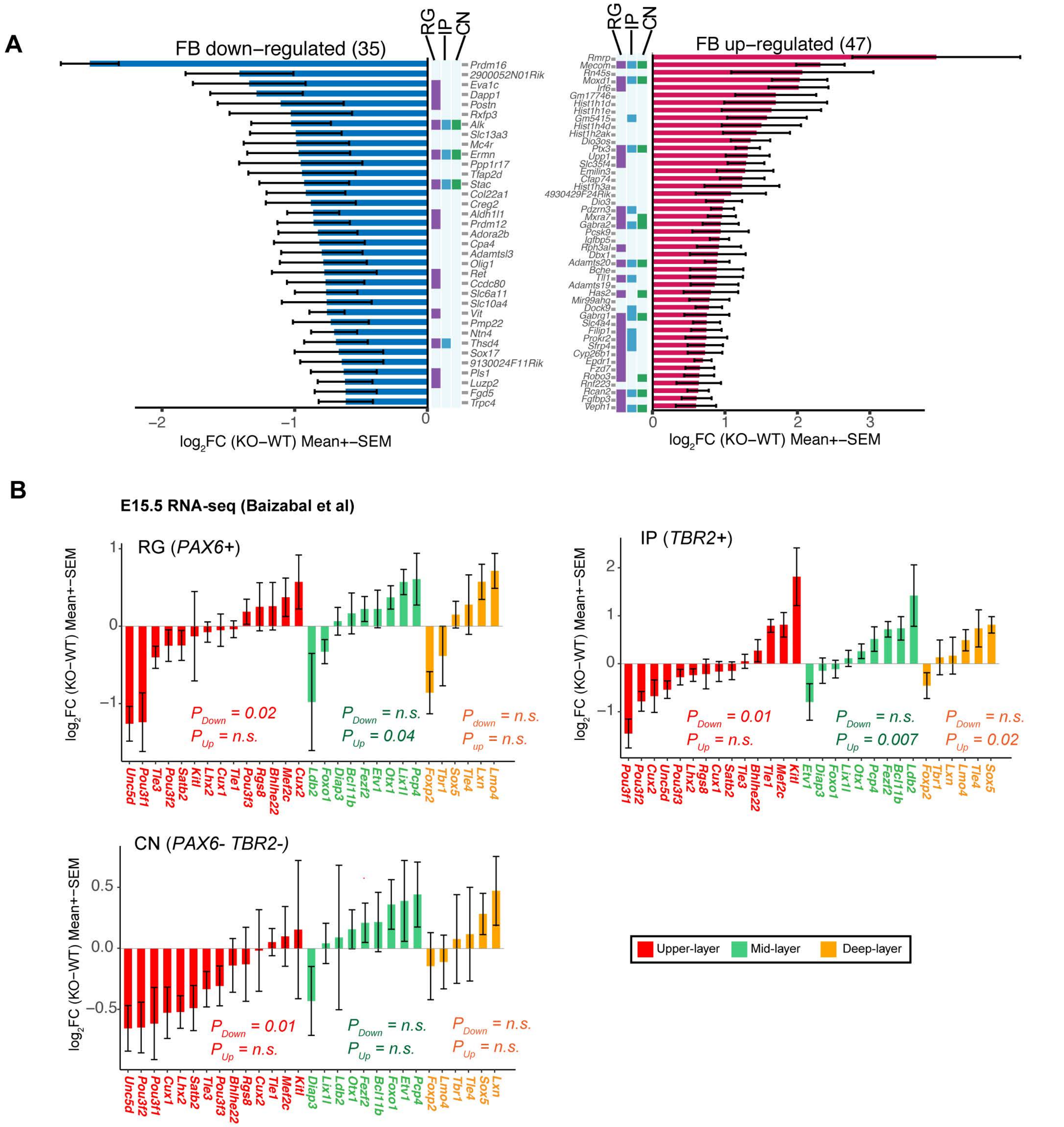


**Figure S4: Top de-regulated genes in the mutant forebrains from RNA-seq analysis**

**(A)** Bar graphs showing down-regulated genes in blue and up-regulated genes in red. Cutoff: $Log_2FC$ (KO-WT) < 0.05. Purple, blue and green squeals indicating genes that also changed expression in RG, IP and CN respectively from Baizabal et al, 2018. **(B)** Fold-changes of upper-layer, mid-layer, and deep-layer markers are shown for E15.5 RG, IP and CN (Baizabal et al). A gene set test was performed to test whether the layer markers are significantly changed as a set. The upper layer markers are significantly

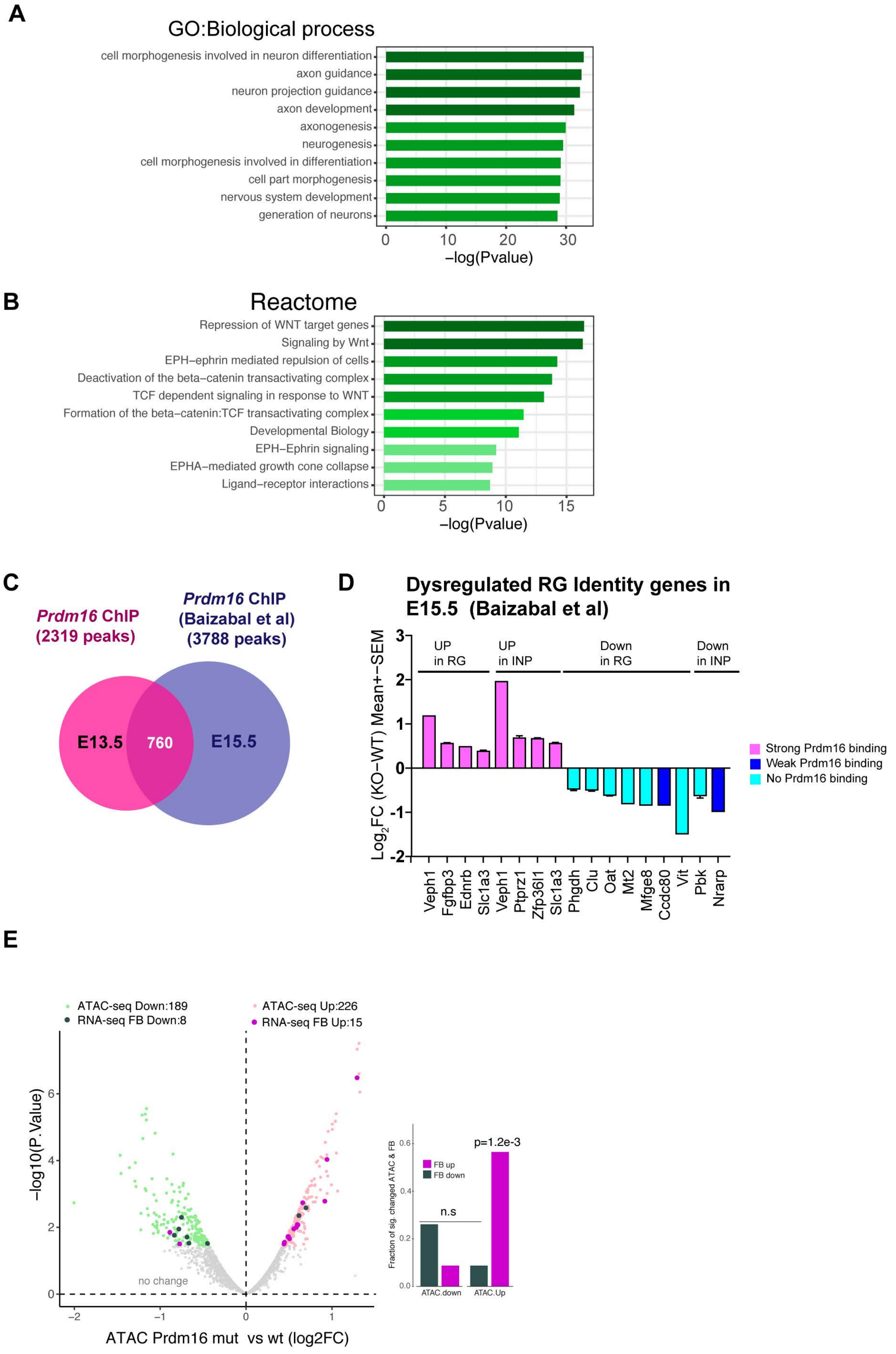
A
GO:Biological process
cell morphogenesis involved in neuron differentiation
axon guidance
neuron projection guidance
axon development
axonogenesis
neurogenesis
cell morphogenesis involved in differentiation
cell part morphogenesis
nervous system development
generation of neurons
0 10 20 30
−log(Pvalue)
B
Reactome
Repression of WNT target genes
Signaling by Wnt
EPH−ephrin mediated repulsion of cells
Deactivation of the beta−catenin transactivating complex
TCF dependent signaling in response to WNT
Formation of the beta−catenin:TCF transactivating complex
Developmental Biology
EPH−Ephrin signaling
EPHA−mediated growth cone collapse
Ligand−receptor interactions
0 5 10 15
−log(Pvalue)
C
Prdm16 ChIP (2319 peaks)
Prdm16 ChIP (Baizabal et al) (3788 peaks)
E13.5
760
E15.5
D
Dysregulated RG Identity genes in E15.5 (Baizabal et al)
Log2FC (KO−WT) Mean+−SEM
UP in RG
UP in INP
Down in RG
Down in INP
Veph1
Fgfbp3
Ednrb
Slc1a3
Veph1
Ptprz1
Zfp36l1
Slc1a3
Phgdh
Clu
Oat
Mt2
Mfge8
Ccdc80
Vit
Pbk
Nrarp
Strong Prdm16 binding
Weak Prdm16 binding
No Prdm16 binding
E
ATAC-seq Down:189
RNA-seq FB Down:8
ATAC-seq Up:226
RNA-seq FB Up:15
−log10(P.Value)
no change
ATAC Prdm16 mut vs wt (log2FC)
Fraction of sig. changed ATAC & FB
FB up
FB down
n.s
p=1.2e-3
ATAC.down
ATAC.Up

**Figure S5:** Prdm16 peak functional enrichment are for **(A)** nervous system development in Gene Ontology (GO) “Biological Process”, and **(B)** migration signaling pathways such as Eph signaling and semaphorin signaling and for RG function such as repression of WNT target genes in “Reactome”. **(C)** Comparison of Prdm16 targets in E13.5 whole brain (this study) and in E15.5 cortex (Baizabal et al). **(D)** A subset of the RG core identity genes bound by PRDM16 shows up-regulation in Prdm16 mutant RG or IP while those weakly (low peak score) or not bound by Prdm16 show down-regulation in Prdm16 mutant. (**E**) A volcano plot showing significantly increased (light pink) or decreased (light green) ATAC-seq signal in mutant vs control at PRDM16-bound loci (FDR <= 0.2). The associated genes that had expression change in the mutant FB were indicated in purple (up-regulated) or dark-green (down-regulated). The bar plots on the right sides measure the fraction of gene expression change in FB over the significantly changed ATAC signals on the Prdm16-bound peaks, showing the up-regulated genes are associated with increased ATAC-seq signal.

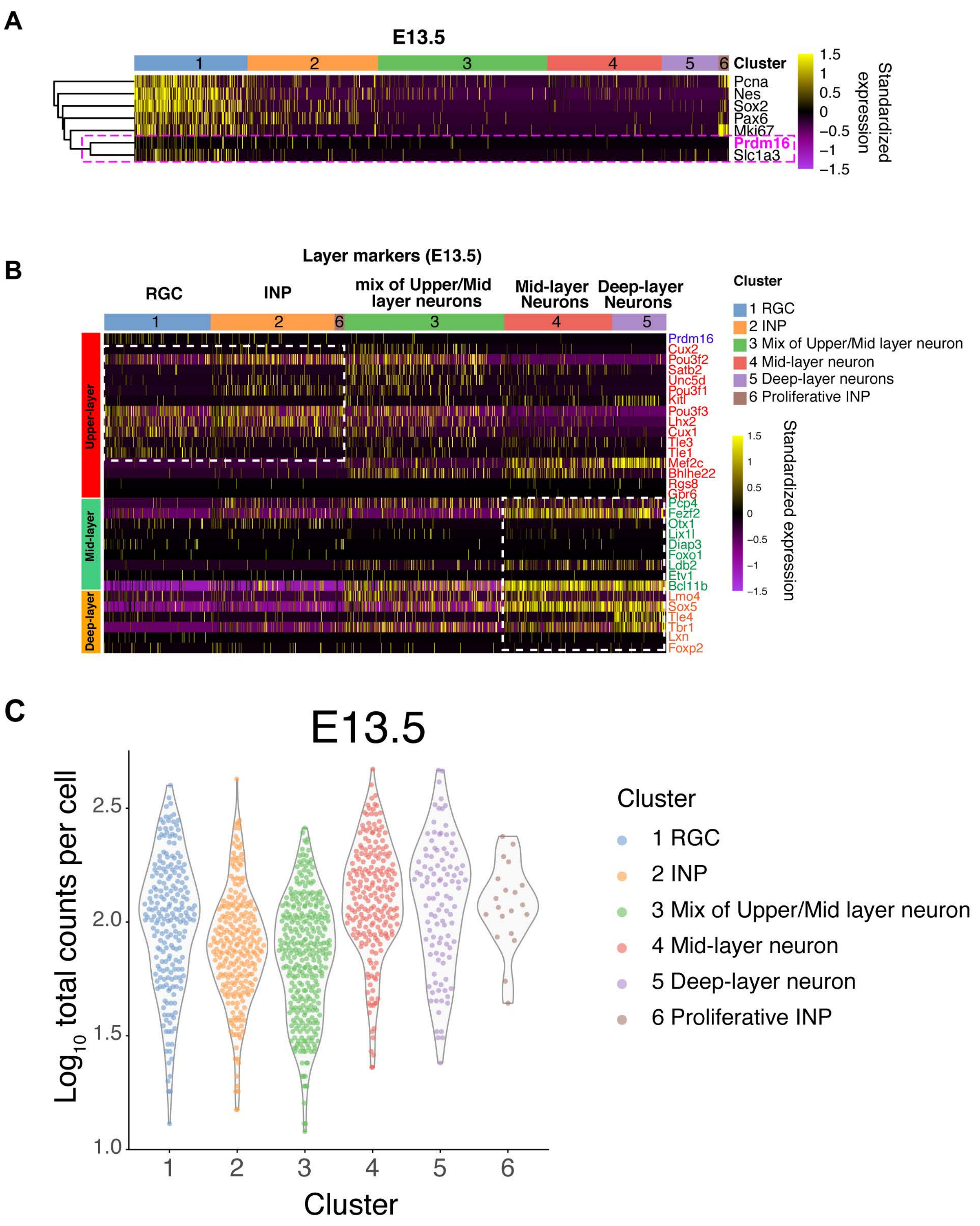


**Figure S6: (A)** Prdm16 is mainly expressed in the RG clusters and is highly correlated with the RG marker Slc1a3 at E13.5. **(B)** Single cell expression for upper-, mid- and deep-layer markers at E13.5. **(C)** Single cell expression for Prdm16 target genes at E13.5. Each dot in the volin represents the Prdm16 target expression (log10 total reads in each cell in each cluster).

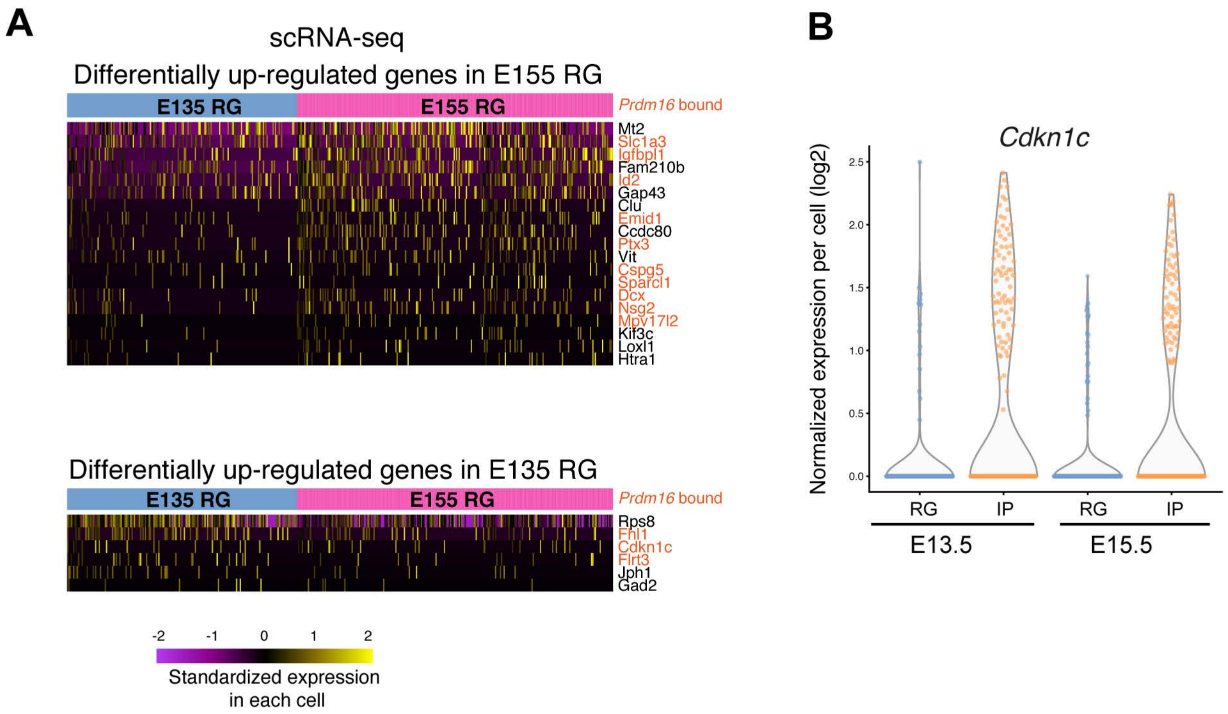


**Figure S7: (A)** Among differently expressed genes between E15.5 and E13.5 RG cells, 24 genes showed most significant changes in *Prdm16* mutant RG (FDR <0.05). The heatmap of these 24 genes in the 13.5 (6 genes) and 15.5 RG (17 genes) scRNA-seq cells are shown. The genes that associate with PRDM16 ChIP peaks are highlighted in orange. (B) A violin plot shows normal expression of *Cdkn1c* in E13.5 and E15.5 RG and IPs.
